# Identifying and removing widespread signal deflections from fMRI data: Rethinking the global signal regression problem

**DOI:** 10.1101/662726

**Authors:** Kevin M. Aquino, Ben D. Fulcher, Linden Parkes, Kristina Sabaroedin, Alex Fornito

## Abstract

One of the most controversial procedures in the analysis of resting-state functional magnetic resonance imaging (rsfMRI) data is global signal regression (GSR): the removal, via linear regression, of the mean signal averaged over the entire brain, from voxel-wise or regional time series. On one hand, the global mean signal contains variance associated with respiratory, scanner-, and motion-related artifacts. Its removal via GSR improves various quality control metrics, enhances the anatomical specificity of functional connectivity patterns, and can increase the behavioural variance explained by such patterns. On the other hand, GSR alters the distribution of regional signal correlations in the brain, can induce artifactual anticorrelations, may remove real neural signal, and can distort case-control comparisons of functional-connectivity measures. Global signal fluctuations can be identified by visualizing a matrix of colour-coded signal intensities, called a carpet plot, in which rows represent voxels and columns represent time. Prior to GSR, large, periodic bands of coherent signal changes that affect most of the brain are often apparent; after GSR, these apparent global changes are greatly diminished. Here, using three independent datasets, we show that reordering carpet plots to emphasize cluster structure in the data reveals a greater diversity of spatially widespread signal deflections (WSDs) than previously thought. Their precise form varies across time and participants and GSR is only effective in removing specific kinds of WSDs. We present an alternative, iterative correction method called Diffuse Cluster Estimation and Regression (DiCER), that identifies representative signals associated with large clusters of coherent voxels. DiCER is more effective than GSR at removing diverse WSDs as visualized in carpet plots, reduces correlations between functional connectivity and head-motion estimates, reduces inter-individual variability in global correlation structure, and results in comparable or improved identification of canonical functional-connectivity networks. All code for implementing DiCER and replicating our results is available at https://github.com/BMHLab/DiCER.

## Introduction

Resting-state functional magnetic resonance imaging (rsfMRI) involves recording spontaneous fluctuations of the blood-oxygenation-level-dependent (BOLD) signal as individuals lie quietly in a scanner without performing an explicit task. It has become a popular and powerful tool for probing brain functional organization in health and disease (1, 2). The dynamics recorded under such conditions are heritable (3, 4); show moderate reliability (5–7); can be used to identify individual participants (8, 9) or discriminate between patient and control groups (2, 10); influence task-evoked activity and behaviour (11, 12); and are spatially organized into functionally related and anatomically connected systems (13, 14). However, despite these and other applications of the technique to address diverse questions in cognitive and clinical neuroscience, there is ongoing debate over how such data should be optimally preprocessed or, more specifically, denoised (5, 15–20).

BOLD signal fluctuations recorded with fMRI contain contributions from numerous physiological and non-physiological sources. Physiological sources include both neuronal and non-neuronal contributions, such as those arising from respiratory and cardiac cycles (21–23). Non-physiological sources include thermal and scanner noise, reconstruction artifacts, and head motion (5, 15, 24, 25). An effective denoising pipeline should isolate the neuronal component of the BOLD signal from other physiological and non-physiological contributions. In task-based fMRI, contrasts between different conditions can be used to ‘subtract out’ most sources of constant (task-uncorrelated) noise. Such contrasts are not possible in typical rsfMRI experiments. Consequently, many denoising approaches have been developed to remove non-neuronal sources of BOLD signal variance for rsfMRI. These include: (i) model-based approaches, which estimate and remove BOLD signal contributions arising from measured peripheral physiological sources (21, 26, 27) or head motion (28, 29); (ii) censoring individual time points contaminated by high motion (30, 31); (iii) data-driven techniques that estimate various noise sources from the data using methods such as independent component analysis (ICA) and principal component analysis (PCA) (32–35); and (iv) the combination of data-driven methods with multi-echo acquisitions, which can be used to reliably separate physiological and non-physiological contributions to the BOLD signal (25, 36, 37).

Although some processing pipelines and techniques can be quite effective in mitigating some of the specific artifacts caused by motion, physiology and scanner issues (5, 15, 25, 33, 35, 36), many are unsuccessful in completely removing prominent and spatially widespread signal fluctuations commonly found in fMRI data (22, 25, 38). These fluctuations are most clearly visualized using a heat map of the voxel × time matrix, a so-called ‘carpet plot’ (or ‘grayplot’), in which the color of each matrix element color represents signal intensity (39). Figure 1 displays signal intensities after a typical preprocessing pipeline that includes motion-related denoising using ICA-AROMA(35), a popular motion-correction strategy, combined with regression of signals estimated from white matter and cerebrospinal fluid (AROMA+2P).

**Fig. 1.**
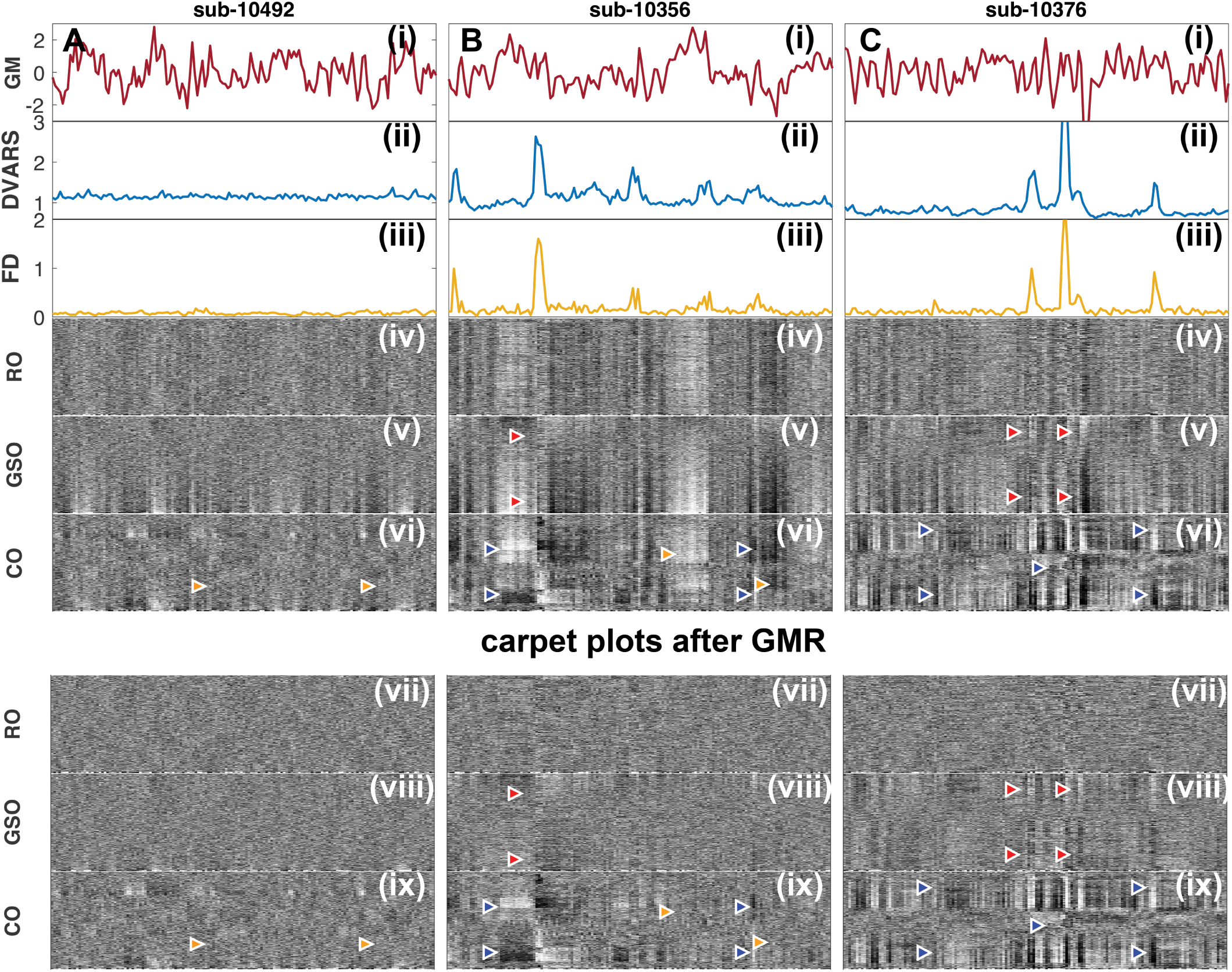
Voxel ordering has a major effect on the visual interpretation of carpet plots. Each column displays a different subject, which represents an example of three heuristic categories of carpet plots found in the UCLA study: **A** low global signal (Subject 10492), **B** prominent single-phase WSDs (Subject 10356), and **C** prominent biphasic WSDs (Subject 10376). The first three rows show: (i) the mean GM signal, (ii) the temporal Derivative of root mean square VARiance over voxelS (DVARS)(77), and (iii) framewise displacement (FD). DVARS and GS are calculated after basic fmriprep preprocessing (before AROMA). The next three rows show carpet plots of the same data plotted using different voxel orderings: (iv) random order (RO) carpet plots; (v) global signal order (GSO) carpet plots; and (vi) cluster order (CO) carpet plots. All heatmaps use the same color scale and present identical data for each individual. The last three rows show the carpet plots after applying GMR (vii),(viii) and (ix) under RO,GSO, and CO respectively. Noise-related structure that exists after GMR is labelled with triangles (colored red for GSO and blue for CO). Green arrows indicate where noise-related structure is successfully removed by GMR.

Despite the denoising, widespread vertical bands of uniformly high or low signal are apparent, consistent with an apparently ‘global’ fluctuation in BOLD signal. Such dominant widespread signal deflections (WSDs) have a dramatic impact on univariate signal properties (e.g., spectral power) and functional connectivity, measured using either traditional (e.g., pairwise Pearson correlations) or time-resolved (e.g., sliding window) approaches (e.g., Zalesky et al. (40)).

A comprehensive study by Power et al. (22) showed that these WSDs are closely tied to non-neuronal influences; principally scanner artifacts, respiratory variations, and head motion (which often coincides with changes in respiration; see also Power et al. (25)). They further showed that explicitly measuring, modeling, and removing motion and respiration-related effects via methods such as RETROICOR (41) does not sufficiently remove WSDs from the data. Instead, Global signal regression (GSR), which uses linear regression to remove the whole-brain average signal from each individual voxel, was the most effective technique for ‘flattening’ the carpet plot; i.e., yielding a carpet plot that visually minimizes WSDs.

GSR is arguably the most controversial preprocessing step used in rsfMRI denoising (16, 42). GSR improves the anatomical specificity of functional connectivity measures by reducing a bias for most voxels to have positively correlated fluctuations (an effect attributable to WSDs) (17). It can mitigate some motion-related confounds in functional connectivity analyses (see (5, 15)) and improve correlations between functional connectivity and behavior (43). GSR also has proven efficacy in removing signal contributions from respiratory and other non-neuronal physiological processes (21, 22). However, GSR has notable drawbacks. GSR mathematically forces the distribution of Pearson correlations to be centered on zero, complicating the interpretation of ‘negative correlations’, which may be introduced artifactually (17, 18, 44). This effect is not simply a relative shift of the original distribution of correlations with positive mean; rather, GSR can alter BOLD signal covariances in a spatially heterogeneous way, depending on the initial size and strength of correlations between clusters of voxels (18, 45, 46). This alteration can lead to spurious differences in case-control comparisons of functional connectivity (5, 18, 45, 47, 48), although the severity of this effect may depend on the dimensionality of the data (49). Furthermore, evidence that the global-mean signal contains neuronal contributions (50–52), some of which are behaviourally relevant (53–55), suggests that GSR may remove relevant signal. Indeed, Glasser et al. (46) found that application of GSR and ICA–FIX (33) to task fMRI data in the HCP led to reduced statistical sensitivity for detecting activations relative to the application of ICA– FIX alone. GSR can also exacerbate the impact of motion on short-range compared to long-range connections (5, 15). A final and under-appreciated limitation of GSR, which we discuss in detail below, is that the estimated ‘global signal’ is often not ‘global’; i.e., it does not reflect a common signal present in all, or even the majority, of the brain’s voxels. Instead, it contains contributions from different subsets of temporally coherent voxels.

These limitations of GSR underscore the need to develop alternative methods for removing WSDs from BOLD data. Glasser et al. (46) recently applied temporal ICA (tICA) to data already denoised with spatial ICA (sICA) (more specifically, the FSL–FIX algorithm; (33)). The tICA was used to separate distinct sources of neuronal and non-neuronal signals with widespread anatomical distributions, thus allowing the selective removal of non-neuronal components from the data. However, a limitation of this approach is that it can only be applied on high spatial and temporal resolution data such as those acquired in the Human Connectome Project (HCP) (see also (19, 20)). An alternative approach used a `Go Decomposition` (56), which isolates low-rank components of the voxel time series and yields comparable performance to GSR (25). However, low-rank components are estimated across all voxel time series – a constraint that implicitly assumes that the global signal is indeed global (i.e., a common signal present across most voxels). As we show in below in see Sec. I, this assumption is only valid in certain cases. Lastly, Erdoğan et al. (57) present a method that shifts the estimated global signal in time and performs voxel-specific regression with the most correlated shift at each voxel. This approach was shown to improve estimation of resting-state networks, but it is also applied at a global scale.

In this article, we present a new data-driven approach for identifying and removing WSDs from rsfMRI data. Section I motivates our method with a detailed examination of how WSDs manifest across time and different individuals. We show that the diversity of WSDs that occur in rsfMRI data can be visualized effectively by reordering the rows of the carpet plot to emphasize subject-specific spatial patterning of the data. This structure is hidden in conventional carpet plots, which can be misleading as a tool for evaluating the efficacy of denoising methods. In Section II, we introduce a new, iterative cluster-based method for removing WSDs which assumes that synchronized signals that extend broadly across the whole brain are either artifactual or not important for understanding distributed information processing. Our algorithm, called Diffuse Cluster Estimation and Regression, DiCER, iteratively searches for different sources of WSDs using the clustering algorithm `dbscan`(58), regressing out the contribution of any WSD estimated at each iteration. The approach is more flexible than GSR for identifying artifacts with irregular signal fluctuations (e.g., sharp spikes or biphasic signal deflections) and, unlike GSR, DiCER only applies correction when it finds evidence of WSDs. Critically, DiCER does not force a particular correlation distribution on the denoised data, and so it does not introduce anticorrelations by construction (a noted limitation of GSR (17, 44)). Lastly, in Section III, we compare the performance of DiCER to GSR in: (i) removing WSDs (through inspection of carpet plots); (ii) mitigating correlations between functional connectivity and head motion; and (iii) identifying anatomically plausible functional-connectivity networks. All code for implementing DiCER and reproducing our results is available at https://github.com/BMHLab/DiCER.

## Section I: Visualizing voxel time series

### Imaging methods

#### fMRI data

We used open source rsfMRI datasets processed using transparent pipelines from `fmriprep` (59). We focus in particular on data from the healthy controls of the UCLA Consortium for Neuropsychiatric Phenomics LA5c Study (60) (v00016 openneuro.org/datasets/ds000030/), and use the Beijing-Zang dataset (fcon_1000.projects.nitrc.org/fcpClassic/FcpTable.html) and multi-echo Cambridge dataset (25) (v00002 openneuro.org/datasets/ds000258/ we focused on the 2nd echo (TE = 32 ms)) for replication.

The scanning parameters for these three datasets are described in (59), http://fcon_1000.projects.nitrc.org/fcpClassic/FcpTable.html and (25), respectively; a brief summary is in Table 1. For clarity, we separate our preprocessing methods into two sections: (i) steps carried out with `fmriprep` (59); and (ii) denoising steps performed subsequently.

**Table 1.** Summary of acquisition parameters for the functional MRI and structural MRI used in this study. Note: not all essential parameters were reported in the open-source repositories, we list those that were reported.

| Dataset | BOLD parameters | Volumes | Subjects | Notes |
| --- | --- | --- | --- | --- |
| UCLA LA5c Study (60) | TE = 30 ms, 3 mm Inplane resolution, 34 slices with 4 mm slice thickness, FA= 90, FOV= 192 mm, matrix = 64 × 64, Oblique and interleaved Gradient echo EPI sequences, TR = 2 s | 152 | 121 | We focused on the healthy controls (from the original sample of 270 people), that included subjects aged 21–50. |
| Beijing-Zang | TE= 30 ms, 3.125 mm, Inplane resolution, 33 slices with 3.6 mm, slice thickness, FA= 90, FOV= 200 mm, matrix = 64 × 64 | 225 | 192 | Subjects were healthy controls. |
| Cambridge Multiecho (25) | TE = 12, 28, 44, 60 ms, 3.75 mm Inplane resolution, 32 slices with 4.4 mm slices thickness FA = 78, FOV = 40 mm, interleaved slice acquisition with 10% gap Gradient echo EPI, iPAT = 3, TR = 2.47 s | 139 | 89 | Subjects were healthy controls. We only analyzed the second echo TE = 28 ms. |

#### fmriprep workflow

Results included in this manuscript come from preprocessing performed using `fmriprep` v1.1.1 (59, 61), a Nipype-based tool (62, 63). Each T1-weighted (T1w) volume was corrected for intensity non-uniformity using `N4BiasFieldCorrection` v2.1.0 (64) and skull-stripped using `antsBrainExtraction.sh` v2.1.0 (using the NKI template). Brain surfaces were re-constructed using `recon-all` from FreeSurfer v6.0.1 (65). A brain mask was estimated with FreeSurfer, which was refined using an atlas based brain mask, similar to that within Klein et al. (66). Spatial normalization to the ICBM 152 Nonlinear Asymmetrical template version 2009c (67) was performed through nonlinear registration with the `antsRegistration` tool of ANTs v2.1.0 (68), using brain-extracted versions of both T1w volume and template. Brain-tissue segmentation of cerebrospinal fluid (CSF), white-matter (WM) and gray-matter (GM) was performed on the brain-extracted T1w using `fast` (69) (FSL v5.0.9).

Functional data were slice-time corrected using `3dTshift` from AFNI v16.2.07 (70) and realigned to a mean reference image using `mcflirt` (71). ‘Fieldmap-less’ distortion correction was performed by co-registering the functional image to the intensity-inverted T1w image (72, 73) constrained with an EPI distortion atlas presented in Treiber et al. (74) and implemented with `antsRegistration` (ANTs). This was followed by co-registration to the corresponding T1w using boundary-based registration (75) with nine degrees of freedom (`bbregister` within FreeSurfer v6.0.1). The motion-correcting transformations, field-distortion-correcting warp, BOLD-to-T1w transformation, and T1w-to-template (MNI) warp were concatenated and applied in a single step using `antsApplyTransforms` (ANTs v2.1.0) using Lanczos interpolation (76). Framewise displacement was calculated for each functional run using the implementation of Nipype, based on the formulation by Power et al. (77). ICA-based Automatic Removal Of Motion Artifacts (AROMA) was used to generate noise regressors for use in the non-aggressive variant of the method (35).

For more details of the `fmriprep` pipeline we refer the reader to https://fmriprep.readthedocs.io/en/latest/workflows.html and the accompanying paper by Esteban et al. (59).

#### Post-fmriprep processing

For transparency, the methods for post-`fmriprep` processing are described in terms of the outputs from `fmriprep` v1.1.1, which detail the steps within our processing tools located at https://github.com/BMHLab/DiCER.git. Functional MRI data are analyzed within the MNI 152 Asymmetric 2009c space, which has been resampled to the native BOLD imaging dimensions, labeled by the tag `bold_space-MNI152NLin2009cAsym`, and we resampled any remaining anatomical masks/images to this space (including those that were not automatically resampled in the `fmriprep` workflow).

Following `fmriprep`, the automatically labeled noise and BOLD ICA components from ICA-AROMA (described above) were used to perform a non-aggressive variant of ICA-AROMA on the unsmoothed outputs of `fmriprep`, labeled by the suffix `preproc.nii.gz`. Regressors were calculated on the spatially smoothed variant (as described within `fmriprep`) of `preproc.nii.gz` and then applied to the unsmoothed preprocessed file.

In what follows, we restrict our analysis and visualizations to gray-matter voxels (GM) for two reasons: (i) we are primarily interested in networks within GM; and (ii) the global-mean signal and mean gray-matter signal are highly correlated (16, 46). To minimize partial-volume effects, our analysis was restricted to voxels contained within the GM probability masks thresholded at > 50% probability. We also excluded voxels with signal intensities that were below 70% of the mean fMRI signal intensity to avoid contamination by voxels with low signal plagued by susceptibility and partial-volume effects.

#### Standard denoising

Following ICA-AROMA, we extracted mean time courses from eroded masks of the WM and CSF. The masks were generated by following Parkes et al. (5) and Power et al. (22), where CSF and WM ROIs were created from tissue probability maps in `fmriprep`. We eroded the WM mask five times and the CSF mask once. Erosion is crucial to avoid partial-volume effects from gray matter, which inflates the correlation between WM/CSF estimates and the global-mean signal (5, 77). We extracted these signals from the AROMA-denoised data, as performed in Pruim et al. (35). Traditionally, the global-mean signal has been estimated by taking the mean BOLD time series across the entire brain mask, including WM and CSF. It has been shown that gray matter makes the strongest contribution to this global mean (22, 46), and that WM and CSF contributions beyond partial-volume effects are negligible (77). We thus use the mean gray-matter signal as a proxy for the global signal, and refer to regression of this signal from fMRI data as gray-matter regression (GMR) for clarity, as this better captures our method than global-signal regression (GSR).

Using the noise-signal estimates, we perform two variants of noise correction: (i) regression with the WM and CSF physiological signals, denoted as ‘+2P’; and (ii) regression with WM, CSF and GM signals, denoted as ‘+2P+GMR’. These models were applied after ICA-AROMA denoising in a single step using ordinary least squares regression implemented in `fsl_regfilt`. The data were then detrended with a 2nd order polynomial and high-pass filtered at 0.005 Hz using AFNI’s `3dTproject`. This procedure resulted in two datasets for each person, labeled ‘ICA-AROMA+2P’, and ‘ICA-AROMA+2P+GMR’.

#### Carpet plots and voxel ordering

The visualization of all voxel time series as a carpet plot (or grayplot), is a very useful tool in detecting and understanding artifacts (39) and hence evaluating noise-correction techniques (77). The carpet plot is a heatmap visualization of the voxel × time matrix, *X* (*V* × *T* for *V* voxels and *T* time points), usually after *z*-scoring each voxel’s time series (rows of *X*), resulting in the normalized matrix 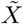 used here. To prevent outliers from dominating the color-scale limits and obscuring the interesting signal, we fixed these limits to the range [− 1.2, 1.2]. Plotting independently measured time series, e.g., of head-motion or respiration, above the carpet plot allows relationships between BOLD signals and non-neuronal time courses to be visualized straightforwardly. Here we focus on individual voxels as spatial elements, but note that carpet plots can also be constructed at coarse-grained levels, for example, grouping voxels as regions of interest.

Historically, voxels in carpet plots are ordered somewhat arbitrarily, by the numerical unfolding of the voxel mask into a vector (which is not a standardized procedure across different software packages), or the ordering of regions in a whole-brain parcellation (which varies depending on the parcellation used). We denote this ordering of voxels as ‘random ordering’ (RO). RO carpet plots reveal artifacts and prominent, whole-brain signals but can obscure more complicated, non-global structure (as we show below). Here we compare two alternative orderings that better reveal spatially structured patterns of signal fluctuations. The first ordering scheme is ‘gray-matter signal ordering’ (GSO), which orders voxels according to their Pearson correlation to the mean GM signal. By placing the voxels with the most positive correlations to the mean GM signal at the top of the plot (and the most strongly negatively correlated voxels are at the bottom), GSO carpet plots facilitate visual inspection of the spatial distribution of a given WSD. The second ordering scheme is ‘cluster-similarity ordering’ (CO), which reorders voxels such that those with similar BOLD dynamics are placed close to one another. We computed this ordering using hierarchical average linkage clustering on Euclidean distances between pairs of *z*-scored time series. CO carpet plots offer a more comprehensive picture of the diversity and spatial extent of WSDs.

#### Visualization results

To investigate the effect of voxel ordering of the carpet plot in revealing large-scale structure in rsfMRI BOLD data, we plotted RO, GSO, and CO carpet plots for three representative individuals in Fig. 1. Data are shown after applying ICA-AROMA+2P (upper plots (iv)– (vi)) and after subsequently applying GMR (lower plots (vii)–(ix)). Note that throughout this paper, voxel orderings are determined from the ICA-AROMA+2P data, and preserved in subsequent visualizations (after a correction method has been applied). This choice is to emphasize signal artifacts after minimal de-noising, thus allowing an evaluation of the efficacy of de-noising techniques at a subject-specific level. In this section, we illustrate our main points using three exemplary subjects, shown in Fig. 1. Additional examples are presented in Fig S1–S3. A full report of similar plots for all individuals across the three datasets considered here (402 subjects in total) can be found at https://bmhlab.github.io/DiCER_results/.

#### WSDs in RO carpet plots

Conventional RO carpet plots, shown in Figs 1A(iv), B(iv), C(iv), reveal large WSDs in all participants. These WSDs appear to be ‘global’, i.e., are present in the vast majority of voxels. Some of these WSDs are tied to head motion, quantified as FD (Figs 1A(iii), B(iii), C(iii)) spikes, whereas other WSDs are not so clearly tied to FD and are likely attributable to respiratory fluctuations (22). Comparing to the carpet plots after applying GMR (Figs 1A(vii), B(vii), C(vii)), prominent WSDs are no longer present, suggesting that they have been successfully removed by GMR (77).

#### WSDs in GSO carpet plots

By construction, GSO carpet plots reveal a gradient of increasing global signal contribuion down the vertical axis, shown in Figs 1A(v),B(v),C(v). For Subject 10492, the GSO plot reveals a consistent common signal in only a small percentage of voxels, which is weakly correlated with FD fluctuations. WSDs are not as pronounced relative to the other two subjects. For Subject 10356, WSDs are visually apparent throughout the vast majority of voxels and are heavily tied to large FD spikes. For Subject 10376, WSDs lead to signal increases in some voxels and signal decreases in other voxels. We thus see a positive correlation with the mean GM signal for some voxels and a negative correlation for others. Some, but not all, of these biphasic WSDs are tied to obvious FD spikes.

The three individuals depicted in Fig. 1 represent exemplars of the different types of WSDs apparent across the individuals in the three datasets analyzed here. That is, we find that individuals either show no or limited WSDs (like Subject 10492), coherent monophasic and (nearly) global WSDs (like Subject 10356), or complex biphasic WSDs (like Subject 10376). To investigate the extent to which these three categories are present across all subjects in the UCLA cohort, we quantified the relationship between voxels in the top half of GSO carpet plots (voxels with below-median correlation to the mean GM signal) and the bottom half (voxels with above-median correlation to the mean GM signal), by computing the correlation between the mean of all below-median voxels (BM) and the mean of all above-median voxels (AM). The separation of voxels between the median is illustrated in Fig. 2A. We find that for subjects like Subject 10492, where WSDs are only present in a minority of all voxels, AM and BM are not strongly correlated, *ρ* = −0.25 (Fig. 2B); for subjects like Subject 10356, where a single WSD persists across the vast majority of voxels, AM and BM are more strongly correlated, *ρ* = 0.32 (Fig. 2C); and for subjects like Subject 10376, where a WSD affects different voxels with different polarities, AM and BM are anticorrelated, *ρ* = −0.48 (Fig. 2D).

**Fig. 2.**
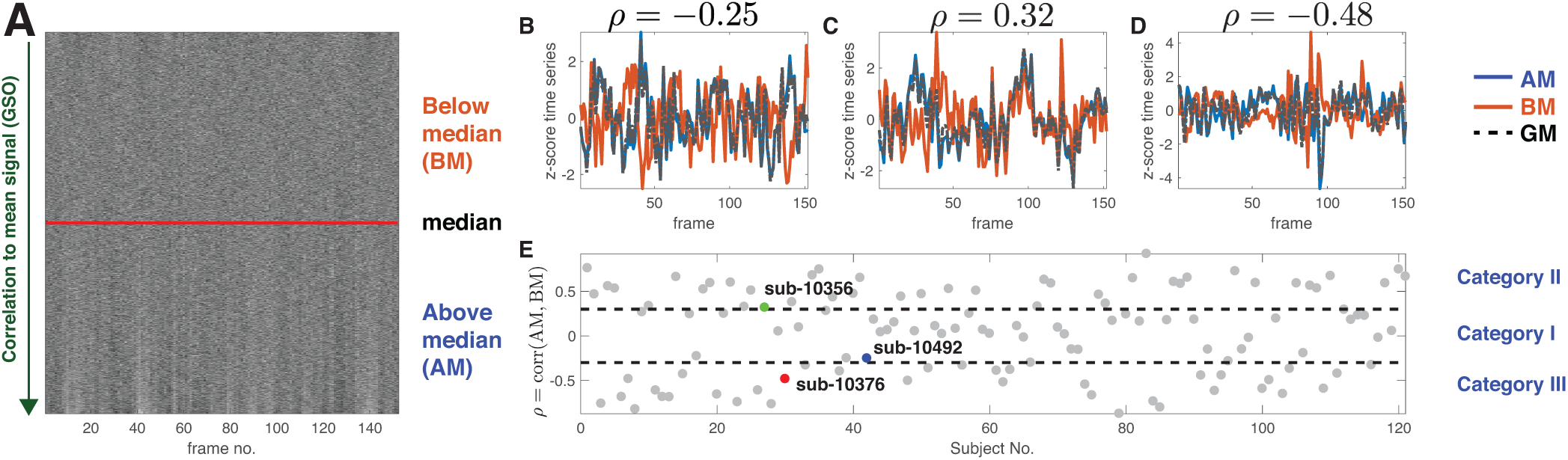
Categorizing carpet plots by their correlation to the mean signal. **A** GSO carpet plot for Subject 10492. In this ordering, the middle of the carpet plot is the median of the correlation of every voxel with the mean signal as indicated by the red line. Voxels below the median (BM) are at the top of the carpet plot and voxels above the median (AM) are shown on the top of the carpet plot. **B**–**D** Signals averaged over BM, AM, and over all gray matter voxels (GM) coloured in orange, blue and dashed-black respectively. The mean signals are calculated from subjects Subject 10492 in **B**, Subject 10356 in **C**, and Subject 10376 in **D** respectively. **E** Correlation of BM with AM, *ρ* = corr(BM,AM), for all subjects in the UCLA cohort. Dashed lines represent the rough classification of carpet plots into three categories: Category I with *ρ ≤* 0.3; Category II with *ρ >* +0.3; and Category III with |*ρ*| < −0.3.

We use the heuristic, *ρ* =corr(BM,AM), to categorize all subjects in the UCLA cohort into one of three categories that correspond to the three qualitative behaviors observed above. As shown in Fig. 2E, we set thresholds on *ρ*, at ±0.3, with Category I subjects exhibiting low |*ρ* | and thus limited evidence for prominent WSDs; Category II subjects exhibiting high *ρ*, thus often expressing coherent and nearly global WSDs; and Category III subjects exhibiting high negative *ρ* consistent with largely biphasic WSDs. This categorization is independent of motion, as neither *ρ* or |*ρ* | are not associated with mean FD across subjects (*r* = −0.08, and *r* = 0.1 respectively). In the UCLA cohort, there are 43 subjects in Category I, 41 in Category II, and 37 in Category III. Note that we use *ρ* here as a simple heuristic for understanding the diversity of WSDs expressed by individuals. We do not suggest that *ρ* supports a natural partition of individuals (indeed, the distribution of *ρ* is continuous, not clustered), nor that every individual fits unambiguously into one of these categories (e.g., some individuals display a mix of monophasic and biphasic WSDs; e.g., Subject 10356 in Figure 1).

Due to the varying distribution of voxel-wise correlations to the mean GM signal across the three categories, GMR has varied effects on the data. For Category I subjects like Subject 10492 (Fig. 1A, cf. Fig. S1), the average of all voxels captures synchronized structure present in the WSD that constructively contributes to the mean while diminishing un-synchronized fluctuations that, on average, cancel each other out. Following GMR, the carpet plot appears ‘flattened’ in RO and GSO carpet plots (Figs 1A(vii), A(viii)).

For Category II subjects like Subject 10356 (Fig. 1B, cf. Fig. S2), the common WSD is represented across the majority of voxels (Figs 1B(iv),B(v)) and is effectively captured in the global average (Figs 1B(i)). GMR again appears to be relatively successful in removing this component from the carpet plots under RO and GSO carpet plots (Figs 1B(vii),B(viii)). However, some WSDs are visible for Subject 10356 under GSO (red arrows in Fig. 1B(viii)), which occur at the time of a large movement spike. These WSDs appear to have a biphasic pattern, in which subsets of voxels show time-locked signal deflections in opposite directions.

Biphasic WSDs are prominent in Category III subjects like Subject 10376 (Fig. 1C, cf. Fig. S3). GMR has limited success in removing WSDs from these subjects because the mean does not adequately capture the global behavior of the brain when voxels show synchronized signal deflections in opposite directions. For example, at the time of major movement events (shown as red arrows in Fig. 1), close to half of the voxels deflect positively while the other half deflect negatively with a similar magnitude, resulting in a near-zero global mean and thus an inability of GMR to correct for the movement event. This can be seen in GSO carpet plots as ‘biphasic’ WSDs after the application of GMR (red arrows in Fig. 1C(v),C(viii)), even though the RO carpet plot appears flattened (Fig. 1C(vii)). Thus, RO carpet plots hide large movement-related artifacts that remain after applying GMR, especially for Category III subjects, which make up approximately one third of the UCLA sample.

#### WSDs in CO carpet plots

The simple GSO voxel ordering is highly revealing of the distribution of global signal across GM voxels and demonstrates how some WSDs can be characterized by large subsets of anticorrelated signal changes. CO carpet plots reveal complex WSDs more clearly, as shown in Figs 1A(vi), B(vi), C(vi) (and Figs S1–S3).

In Category I subjects, such as Subject 10492, CO carpet plots reveal multiple clusters of voxels with different characteristic activity patterns (Fig. 1A(vi)), in addition to the prominent WSDs seen in RO and GSO carpet plots (Figs 1A(iv),A(v)). Some of the minor WSDs [yellow arrows in Fig. 1A(vi), A(ix)] are efficiently captured in the mean signal and are hence effectively removed by GMR, resulting in a flattened CO carpet plot (Fig. 1A(ix)).

For Subject 10356, a representative Category II subject, the CO carpet plot more clearly reveals the extent of biphasic WSDs that occur at large movement events (or the respiratory changes that proceed them (22)), shown in Fig. 1B(vi) (blue arrows). The combination of monophasic and biphasic WSDs in Subject 10356 means that GMR successfully removes some WSDs (yellow arrows) but not others (blue arrows) [see Figs 1B(vi), B(ix)]. These biphasic WSDs are more frequent in Category III individuals like Subject 10376. They are clearly visible with CO and are often time-locked to FD spikes.

The inconsistent mixing of positive and negative deflections in response to movement events means that GMR is more successful at removing some WSDs (yellow arrows in Fig. 1), but is very poor at removing others (blue arrows in Fig. 1). GMR is particularly ineffective at removing biphasic WSDs, as these events result in a near-zero mean signal.

In summary, our understanding of the precise form of WSDs, and the success of GMR in removing them, is highly sensitive to the ordering of voxels in a carpet plot. The traditional RO carpet plot can mask more complicated biphasic WSDs that are time-locked to head motion and not effectively removed by GMR. Critically, because biphasic WSDs are often not prominent in RO plots, such plots can be misleading and suggest that no WSDs are present following GMR when, in fact, there may be many (e.g., Subject 10376 in Fig. 1). The CO carpet plots most clearly represent the diversity of WSDs in a fMRI dataset. Therefore, we use these plots in the remainder of this manuscript.

## Section II: Diffuse Cluster Estimation and Regression for fMRI, DiCER

The previous section indicates that WSDs can exhibit a complex, individual-specific spatiotemporal structure that is often poorly corrected by GMR. In this section, we introduce a method to capture and correct the complex WSD structure highlighted in the CO carpet plots introduced above. Our method assumes that WSDs—large groups of voxels exhibiting highly correlated BOLD dynamics across the recording period—are artifactual and should be removed from fMRI data. This assumption is supported by the striking correspondence between large WSDs and movement/physiological events (e.g., Power et al. (22)). A single iteration of our approach, which we call DiCER, is depicted in Fig. 3. Briefly, it involves applying a clustering algorithm, `dbscan`, to estimates the dominant WSD, and then regressing this signal from the fMRI dataset. The steps in this process are described below.

**Fig. 3.**
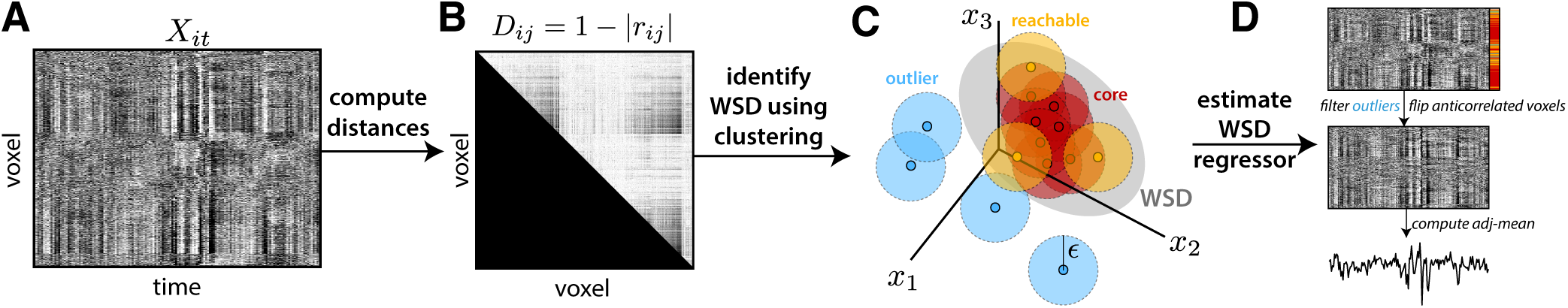
An iteration of DiCER involves using clustering to identify voxels involved in a WSD, and then estimating the WSD regressor as an adjusted mean. We show: **A** CO carpet plot for an example UCLA control participant, Subject 10376. **B** Upper triangle of the pairwise distance matrix, *D*_*ij*_ = 1 − |*r*_*ij*_ |, from low *D*_*ij*_ (black) to high *D*_*ij*_ (white). **C** `dbscan` is used to estimate a diffuse common signal, or WSD, and label the core and reachable voxels that contribute to it. **D** A regressor is estimated from core and reachable voxels, after flipping the sign of voxels that are anticorrelated to the cluster center, as the *adj-mean*. This procedure is repeated until either no WSDs are identified, or a maximum number of iterations, *k*_max_ = 5, is reached.

### Cluster-based WSD estimation

An example fMRI dataset,*X*_*it*_, is plotted as a voxel × time CO carpet plot in Fig. 3A. The first step in DiCER involves identifying WSDs using clustering, which requires us to define a distance measure between pairs of time series. As WSDs can comprise sets of voxels with anticorrelated deflections (Fig. 3A), we treated positively and negatively correlated pairs of voxels equally in our distance measure by taking the magnitude of the absolute Pearson correlation: *D*_*ij*_ = 1 − |*r*_*ij*_|, for a pair of voxel time series, *i* and *j*. As shown in Fig. 3B, this distance measure results in a low distance between pairs of voxels with either strongly anticorrelated or positively correlated time series. In the abstract space defined by *D*_*ij*_, WSDs should manifest as dense regions (clusters), depicted schematically in Fig. 3C. We identify these clusters using the clustering algorithm, `dbscan` (58), which classifies all voxels in a cluster as either: (i) *core* (≥*N*_points_ point within a distance *ϵ*), (ii) *reachable* (within a distance *ϵ* from a core point), or (iii) *outliers* (no core points within a distance *ϵ*). The algorithm parameters, *N*_points_ and *ϵ*, were set with the aim of accurately detecting the types of WSDs characterized above. While `dbscan` is traditionally used to detect compact, highly-similar clusters, here we wish to detect a diffuse, semi-global cluster containing contributions from a large fraction of GM voxels that is consistent with the WSDs characterized above. Through empirical testing across the three datasets considered here, the following parameters performed well for this purpose: *N*_points_ = 0.01*V*, where *V* is the number of voxels (1% of all voxels) and *ϵ* = 0.8 (two voxels, *i, j*, are neighbors if |*r*_*ij*_ |> 0.2). We also enforced a minimum cluster size, such that clusters are only detected that contain at least 10% of all voxels in the core.

Once a diffuse cluster of voxels is identified (Fig. 3C), we next estimate the WSD driving the correlation between these voxels. To this end, we developed an algorithm to compute the adjusted mean signal, *m*^adj^(*t*), at any given time, *t*, which can account for the biphasic signal deflections characterized above. To estimate the adjusted mean of a cluster of voxels, we first define the cluster centroid as the voxel with the minimum *D*_*ij*_ to all other voxels in the cluster, *c*, denoting its timetrace as *v*^(*c*)^(*t*). We then compute 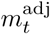 by flipping the sign of voxels that are anticorrelated to the centroid’s time trace:

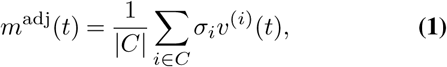

where the sum is taken across all voxels in the cluster set, *C* (of size |*C*|), and the sign indicator σ_*i*_ = 1 if corr[*v*^(*c*)^(*t*), *v*^(*i*)^(*t*)] > 0 and σ_*i*_ = −1 otherwise. The adjusted mean, *m*^adj^(*t*), forms our WSD estimate which is then regressed from the voxel × time data matrix, *X*_*it*_.

### Iterative cluster-based correction of WSDs

To correct for multiple complex WSDs, the above procedure is applied iteratively, with each iteration yielding a progressively corrected fMRI data matrix resulting from the regression of a new WSD estimate, *m*^adj^(*t*). The procedure is terminated when either: (1) no clusters are identified, or (2) a maximum iteration number, *k*_max_ = 5, is reached. Note that DiCER can find zero clusters, and hence apply no correction to the data when it does not find evidence for WSDs, in contrast to GSR, which always regresses out the global mean (and thus centers the distribution of FC values).

For incorporation into a full processing pipeline, the above procedure yields a (possibly empty) set of regressors {*m*_1_, …, *m*_*k*_}, corresponding to the estimated WSDs detected at each of *k* iterations (*k* ≤*k*_max_). Here we used GM voxels to estimate the cluster-based regressors, which were then regressed from all voxels within ICA-AROMA+2P in a single model in FSL using `fsl_regfilt`.

### Visual inspection of DiCER results

To evaluate the effectiveness of DiCER, we applied it to the UCLA, Beijing, and Cambridge cohorts. We took outputs from the ‘ICA-AROMA+2P’ preprocessing stream as input to the cluster-based correction method described above. As shown in Fig. 4, we use Subject 10934 (see Fig. S3), a Category III subject with biphasic WSDs that are not effectively removed by GMR (e.g., Fig S3) as a case study to demonstrate how DiCER iteratively removes complex WSDs.

**Fig. 4.**
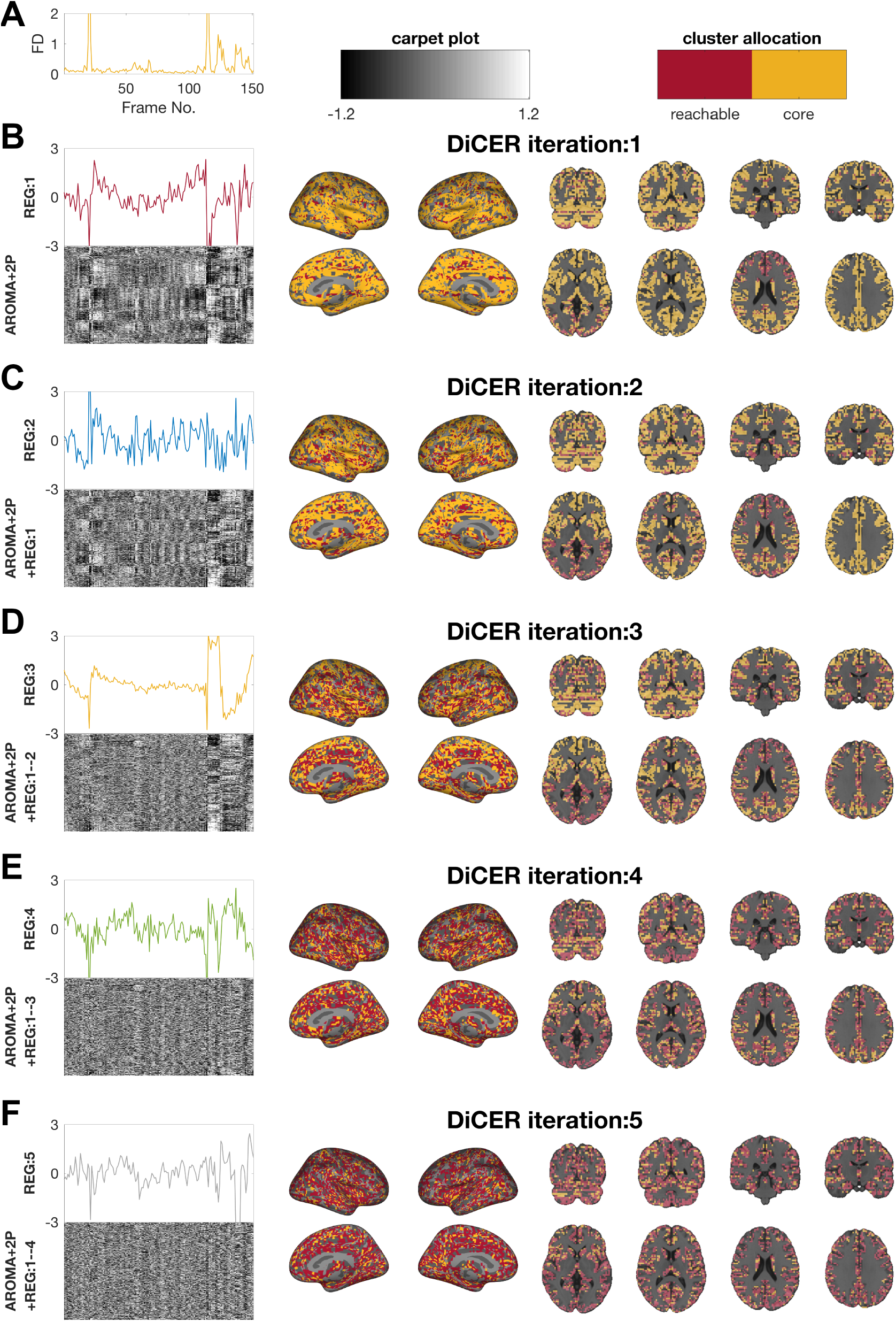
Iterative rsfMRI data correction of Subject 10934 using DiCER. Each row presents an iteration of DiCER, where we plot the DiCER-estimated regressor above the CO carpet plot (from which the regressor was estimated), as well as the distribution of core voxels (yellow) and reachable voxels (red) on cortical surface maps and volume images. The first regressor is estimated from the ICA-AROMA+2P rsfMRI data and removed from each voxel, forming the input for the second iteration, and so on. Signals that have been regressed from the rsfMRI data are labeled on the vertical axis of the carpet plots as ‘+REG:1’, ‘+REG:1–2’, etc.). The FD trace is shown in the top left.

In the first iteration, shown in Fig. 4B, core and reachable voxels have a widespread distribution throughout the brain. There is noticeable banding between core and reachable nodes that is picked up in the coronal slices, the direction orthogonal to the slice acquisition. As the acquisition was interleaved, this banding may caused by the imaging sequence or by slice-time correction. This first regressor identifies large shifts that co-occur with movement events (cf. FD time series in Fig. 4A). Although Regressor 1 includes contributions from the vast majority of voxels, it is only weakly correlated to the global mean signal, *r* = 0.18, due mainly to DiCER’s use of the adjusted mean, which accounts for anticorrelated deviations.

After removing Regressor 1 from every voxel (Fig. 4C), several FD-correlated WSDs remain in the data. Many of these are identified and removed in this iteration (Figs 4D), with **ICA-AROMA+2P+REG:1–2** showing a dramatic reduction in movement-correlated WSDs relative to Fig. 4B. Iteration 3 targets remaining motion-locked WSDs near the beginning and end of the recording (Fig. 4A). The regressor identified at this step also exhibits low-frequency structure which might indicate physiological or head position changes related to arousal levels or sleep [a feature found in temporal ICA of rsfMRI by Glasser et al. (46)], coupled with sudden jerks due to waking. There are fewer core voxels in Iteration 3, and they have a more spatially heterogeneous distribution. Fig 4E shows the successful removal of these WSDs. Signals estimated in remaining iterations are visually related to FD (Fig. 4A) and progressively diminish the remaining residual WSDs (which are weaker, as indicated by the reduction in the number of core voxels).

To investigate the spatial distribution of core voxels across all UCLA participants, we calculated a voxel-wise contribution score that measures the proportion of subjects for which that voxel was labeled as ‘core’ (across all DiCER iterations). As shown in Fig. 5, the contribution score varies spatially across the brain, with a slight preference for insula and medial wall regions. However, the maximum contribution score of any single voxel is 0.25 and we don’t see a strong spatial bias for sources of artifactual signals. Note that all surface maps in this manuscript are displayed on freesurfer’s (http://freesurfer.net) `fsaverage` inflated surfaces, where the projections from MNI to the appropriate surfaces were calculated with registration Fusion (78).

**Fig. 5.**
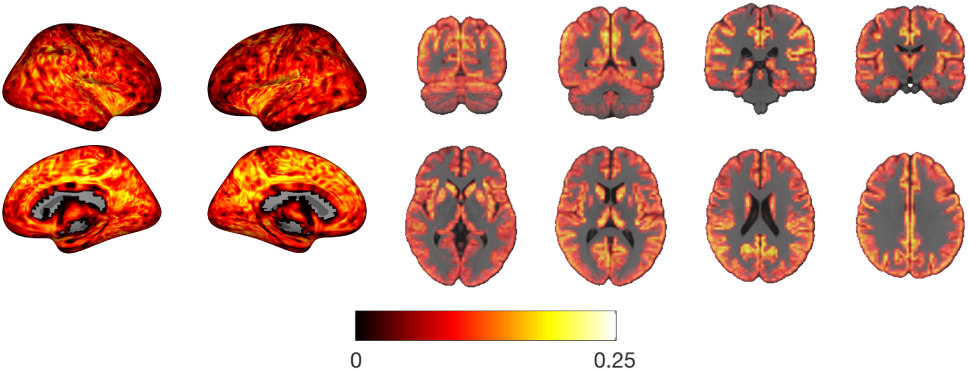
Consistency of core voxel identification across individuals. We plot the spatial map of the participation index, which captures the proportion of subjects for which a given voxel was identified as a core of any nuisance regressor identified by DiCER in the UCLA cohort.

We next demonstrate how DiCER allows better correction of movement-correlated and other WSDs compared to GMR by returning to our three case-study subjects from Fig. 1. CO carpet plots, for all subjects, are shown for (i) uncorrected, (ii) GMR-corrected, and (iii) DiCER-corrected data in Figs 6A, B, and C. For Subject 10492, DiCER finds threeregressors [Figs 6A(vii)–A(ix)] capturing subtle monophasic WSDs and upon regression of these signals the resulting carpet plot is ‘flattened’ [Fig. 6A(iii)]. More examples of Category I subjects are shown in Fig. S4).

**Fig. 6.**
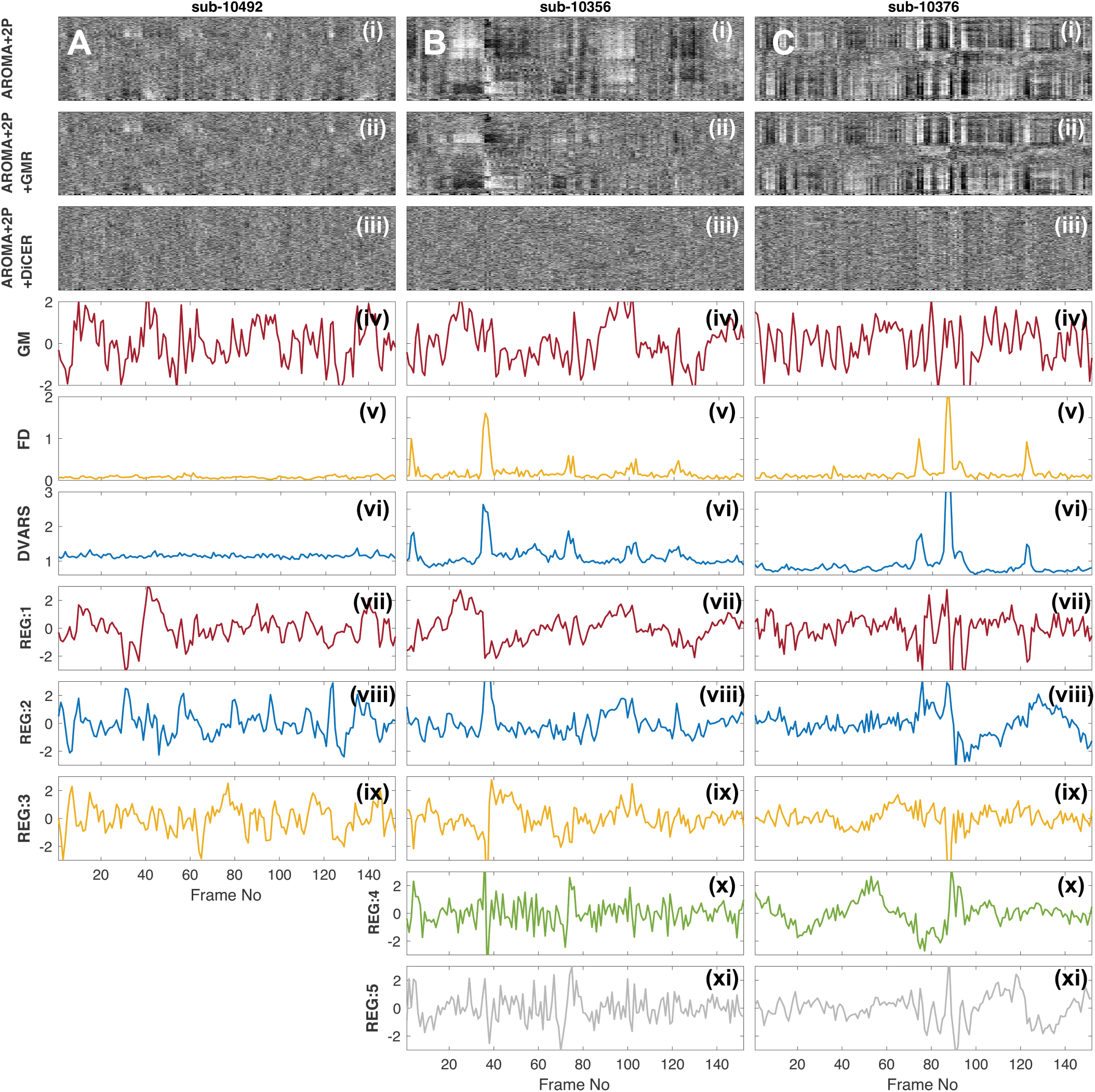
DiCER successfully removes diverse WSDs. Here we show the three subjects from Fig. 1 i.e. Subject 10492, Subject 10356, and Subject 10376 in A,B, and C respectively. The top three rows depict results obtained using three noise-correction schemes: (i) ICA-AROMA+2P, (ii) ICA-AROMA+2P+GMR, and (iii) ICA-AROMA+2P+DiCER, plotted as CO carpet plots. The fourth (iv) row shows the mean grey matter signal, the fifth (v) row shows FD and and the sixth row (vi) shows DVARS. The rows (vii)–(xi) are the regressors estimated by DiCER.

For Subject 10356 [Fig. 6B(ii)], GMR visually fails to remove movement-related WSDs, whereas DiCER successfully removes them through iterative estimation of five regressors [Fig. 6B(iii), regressors shown in Figs 6B(vii)– B(xi)]. Consistent with more complex and varied voxel responses to movement events, DiCER estimates anywhere between 3–5 regressors for Category II subjects. Further examples of Category II subjects are shown in Fig. S5, where DiCER is successful in removing movement-related and other WSDs.

Finally, when encountering the complex biphasic WSDs exhibited by Subject 10376, DiCER estimates regressors that appear motion-related [Regressors 1 and 2 in Figs 6C(vii),C(viii)], as well as further signals that capture additional widespread structure, with both high and low-frequency temporal characteristics around FD spikes [Figs 6C(ix–xi)]. For other Category III subjects (see Fig. S6), DiCER estimates between 3 and 5 regressors.

In some cases DiCER benefits from having multiple regressors, which allow it to capture a greater variety of responses to movement events, and to focus estimation on signals from the subset of voxels that actually display the WSD (rather than always including all voxels, as in GMR). However, even when DiCER estimates a single regressor, it still outperforms GMR, particularly in cases where biphasic WSDs are present. Examples of these cases can be seen in Fig. S7. This is because DiCER flips the sign of anticorrelated voxels to get a better estimate of the WSD.

Detailed reports for all subjects are at: https://bmhlab.github.io/DiCER_results/, which also shows CO carpet plots after applying DiCER correction (`CO_DiCER`).

## Section III: Estimating resting-state networks using DiCER

DiCER visibly removes WSDs from rsfMRI, but does this processing yield sensible, improved estimates of functional connectivity, or does this cleaning over-correct important, neuronally-driven signal in the data? In many cases, WSDs are clearly tied to FD spikes, indicating a clear non-neuronal origin (E.g., Fig 1). In other cases, WSDs have have been tied to respiratory fluctuations (22). However, in the absence of an underlying ground truth, it is difficult to unambiguously evaluate the success of correction methods in removing noise and retaining neuronal signal. In this section we investigate this question in several ways. First, we use commonly used quality-control benchmarks (5, 15) to quantitatively compare the performance of DiCER relative to AROMA+2P and AROMA+2P+GMR. Second, we derive an estimate of the degree to which a carpet plot has been ‘flattened’ and examine how this estimate correlates with head motion before and after DiCER or GMR. Finally, we consider how DiCER influences the properties and identifiability of canonical resting-state functional-connectivity networks.

### Motion dependence

We first consider quality control– functional connectivity correlations (QC–FC), a commonly used benchmark (5, 15, 22), estimated as the cross-subject correlation between FC and mean FD at each connection in a functional connectivity matrix. The QC–FC correlation quantifies the association between inter-individual variance in functional connectivity and gross head motion, as indexed by mean FD (mFD). An efficient denoising method will be less corrupted by motion and hence score lower on this metric. We focus on control individuals from the UCLA sample, excluding high-motion individuals to enable direct comparison to Parkes et al. (5). More specifically, we retained subjects with *mFD* < 0.3 mm, all FD < 5 mm, less than 20% of FD > 0.3 mm and call this the ‘reduced (low motion) UCLA cohort’ which comprised of a subset of 110 subjects. These exclusion criteria were also applied to the Beijing and Cambridge data (shown in the Supplementary material), resulting in cohorts of 125 and 59 individuals, respectively.

We extracted time series for each of 333 regions-of-interest (ROIs) defined by the Gordon parcellation (79) and computed the correlation between each pair of regional time series. For each unique edge in the FC matrix, and each processing pipeline, we computed the cross-subject QC–FC correlation. We summarize the findings as the percentage of *p* < 0.05 (uncorrected) edges in the QC–FC correlation, shown in Fig. 7A,C, and plot the full distribution of QC– FC correlations in Fig. 7B,D. Compared to AROMA+2P, AROMA+2P+GMR shifts the center of the QC–FC distribution nearer to zero, in both UCLA (Fig. 7A) and Beijing (Fig. 7B) cohorts, consistent with the results of Parkes et al. (5) (For results on the Cambridge dataset see Fig. S8). AROMA+2P+GMR also reduces the percentage of *p* < 0.05 (uncorrected) QC–FC correlations from 21.8% to 15.6% in the UCLA cohort (Fig. 7A) and from 20.9% to 6.34% in the Beijing cohort (Fig. 7C). DiCER performs substantially better, narrowing the distribution of near-zero-centered QC– FC correlations and reducing the proportion of significant QC–FC edges with *p* < 0.05 to 9.3% in the UCLA cohort (Fig. 7A) and 5.6% in the Beijing cohort (Fig. 7C).

**Fig. 7.**
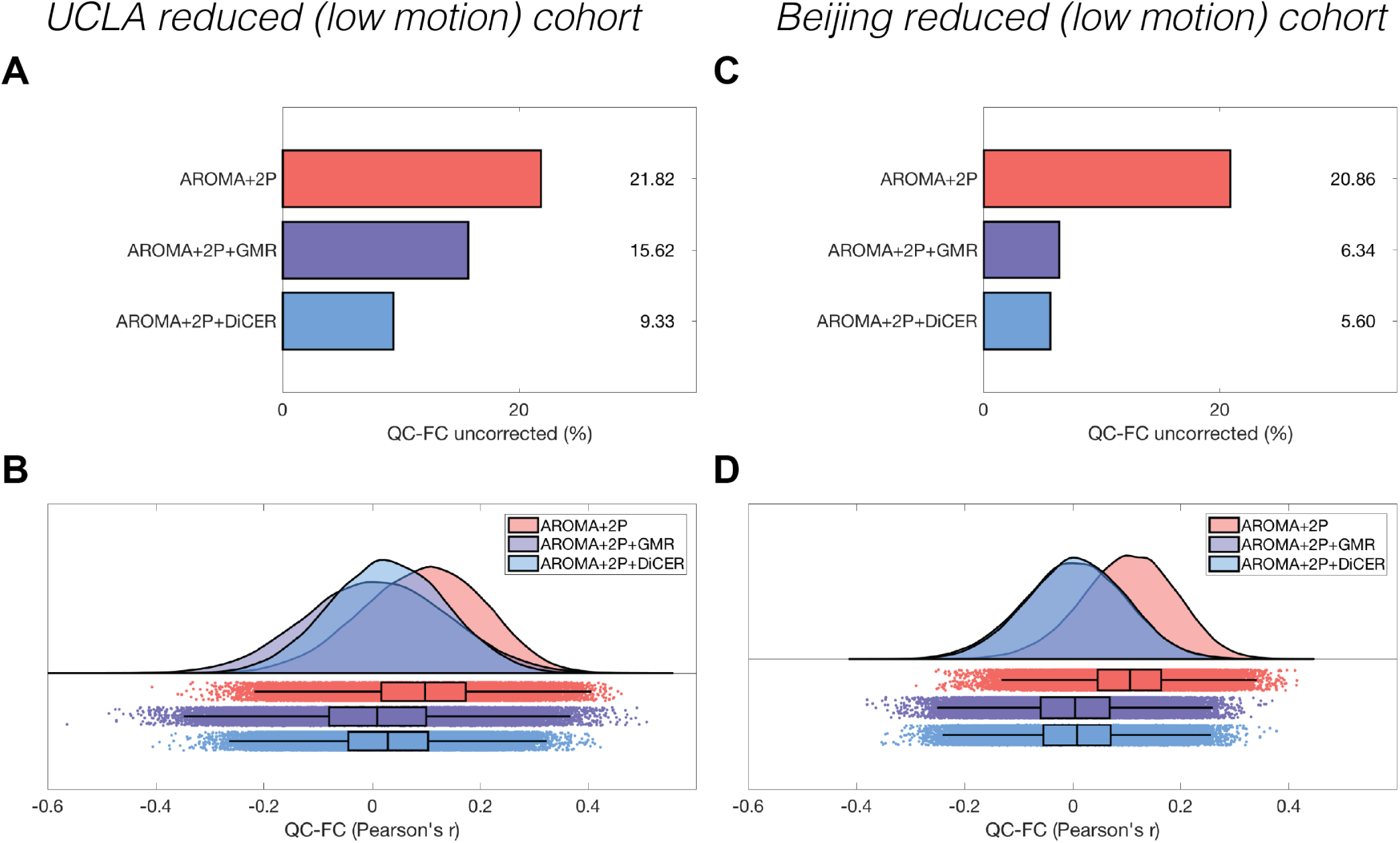
DiCER reduces the correlation between motion (mean FD) and functional connectivity (FC). We compare ICA-AROMA+2P (red), ICA-AROMA+2P+GMR (purple), and ICA-AROMA+2P+DiCER (blue) on the reduced (low-motion) **A, B** UCLA and **C**,**D** Beijing cohorts. **A, C** The proportion of FC values that are correlated to mean FD at a threshold of *p* < 0.05, uncorrected. **B**,**D** The distributions of QC–FC correlations (Pearson’s *r*) across all edges shown as smoothed kernel density estimates at top and boxplots at bottom, with medians and interquartile ranges annotated (80).

Another quantifiable signature of successful rsfMRI denoising is a reduced dependence of QC–FC on the separation distance between pairs of ROIs (5, 24), shown in Fig. 8. Consistent with Parkes et al. (5), GMR reduces the distance dependence of QC–FC in these data in the short, compared to long, connections. However, we also replicate (specifically in the UCLA cohort within Parkes et al. (5)) the finding that GMR can cause a QC–FC pattern that starts positive at close ranges, then turns into a negative correlation before decaying to zero. The distance-dependence of QC–FC following DiCER is reduced with a small increase in mean QC–FC for very short distances, without crossing the zero-axis.

**Fig. 8.**
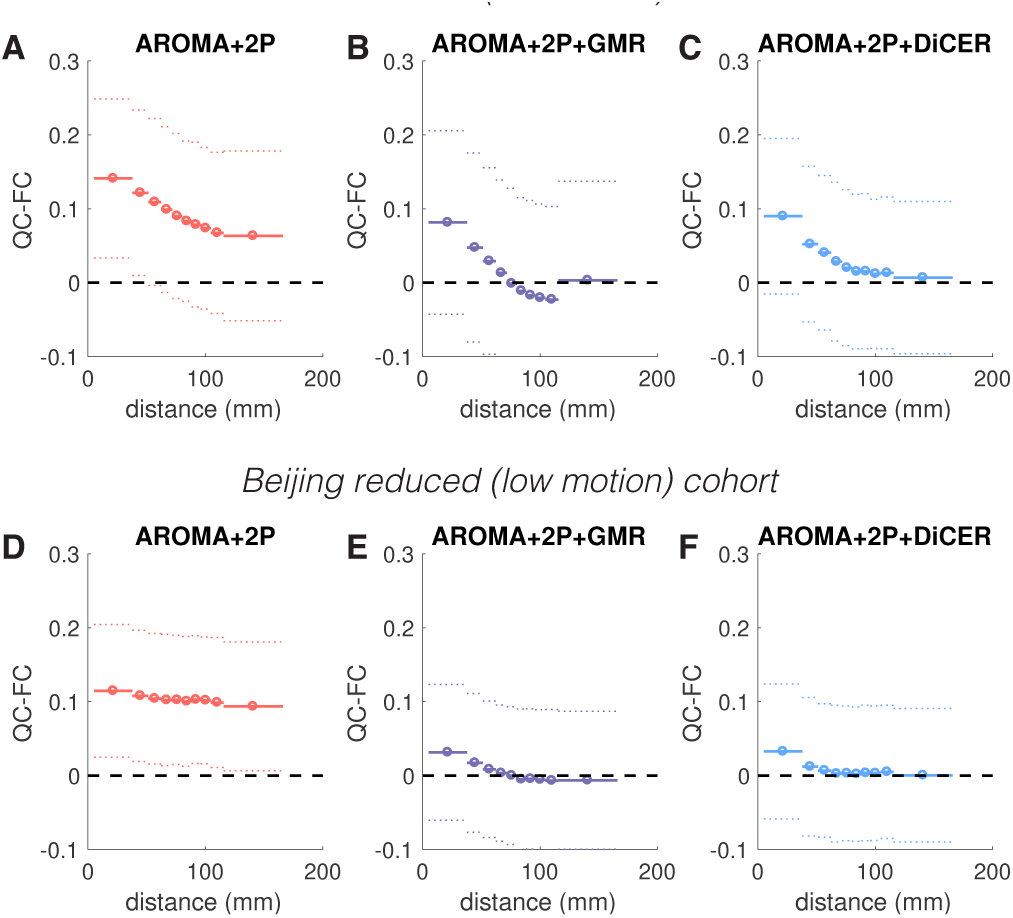
The distance-dependence of QC–FC under different pre-processing pipelines. QC–FC is plotted across ten equiprobable distance bins for the UCL Aand Beijing reduced low motion cohorts, shown as mean (circle and solid horizontal lines) ± standard deviation (dashed horizontal lines), for: **A, D** ICA-AROMA+2P, **B, E** ICA-AROMA+2P+GMR, **C, F** ICA-AROMA+2P+DiCER.

### Towards ‘flat’ carpet plots

In this work, we have used the ‘flat’ appearance of carpet plots to demonstrate that an fMRI dataset does not contain large WSDs, and thus to indicate the success of a given denoising procedure. To quantify the ‘flat-ness’ of a carpet plot, we used the variance explained by the first principal component (PC) of the voxel × time rsfMRI data matrix, which we denote as ‘VE1’. High values of VE1 indicate that considerable fMRI variance can be captured by a single component (PC1), consistent with the presence of dominant WSDs.

The distributions of VE1 across participants in the reduced low-motion UCLA and Beijing cohorts (see above for exclusion criteria) are shown in Fig. 9. For the Cambridge dataset see Fig. S10. DiCER substantially reduces the mean VE1 across participants, whereas GMR has a negligible effect rel-ative to the AROMA+2P pipeline. Note that DiCER has more degrees of freedom than GMR, in that it can remove up to five regressors for each individual. In this sense, we expect the dimensionality of the data to be reduced following DiCER compared to GMR. However, residual WSDs remaining after any of the denoising techniques will lead to the bulk of signal variance loading on to the first PC, and Fig. 9 shows a clear reduction of VE1 following DiCER. DiCER also dramatically reduces the variance of VE1 across individuals, indicating that it is highly consistent in removing WSDs, leaving a similar level of residual coherent activity for all individuals. In contrast, the distribution of VE1 following GMR has an extended tail such that the first PC explains nearly 30% of the variance in some individuals. This large variance in VE1 of GMR-cleaned rsfMRI datasets can be attributed to the diversity of WSDs exhibited by different subjects characterized above in Sec. I. That is, GMR will likely performs well for Category I and II subjects, but not for Category III subjects, or those with a mixture of monophasic and biphasic WSDs. For example the subjects with very high VE1 in the AROMA+2P, and AROMA+2P+GMR pipeline for these two datasets (shown as an asterisk, *, in Figs 9A and B) have large residual WSDs following GMR (see Fig S6 and the online report https://bmhlab.github.io/DiCER_results).

**Fig. 9.**
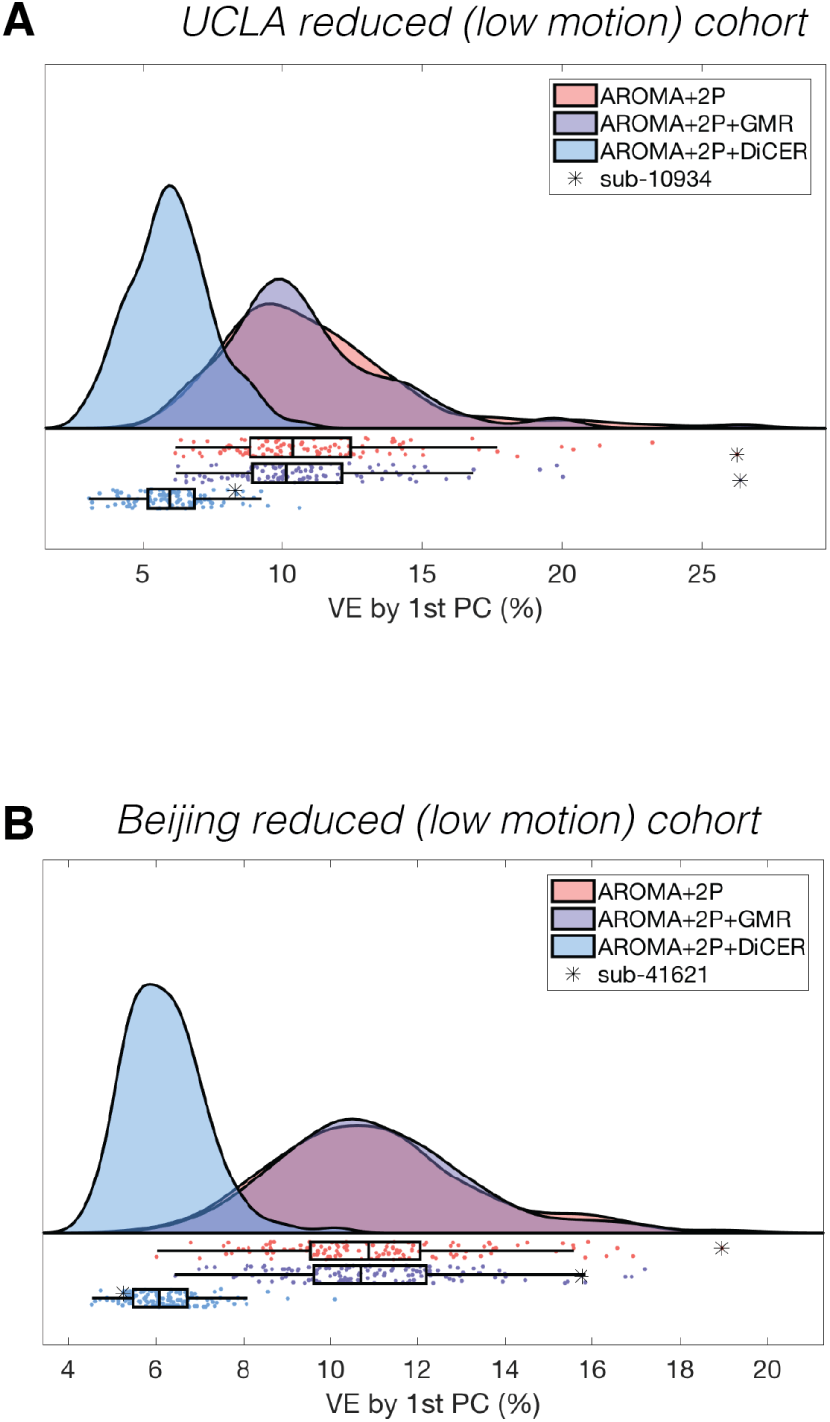
The variance explained (% of total variance) by the first principle component of the carpet plots (VE1) for **A** the reduced (low motion) UCLA, and **B** the reduced (low motion) Beijing cohorts, where the distributions of VE1 are shown as smoothed kernel density estimates at top and boxplots at bottom, with medians and interquartile ranges annotated (80). Asterisks identifying individual subjects for whom GMR did not adequately remove WSDs (see Figs S3, and https://bmhlab.github.io/DiCER_results/BZ/general_report.html).

Note that for subjects in the reduced UCLA cohort, mFD is not strongly correlated with VE1 across all subjects and for all noise-correction techniques (Fig. S11). This indicates that the prominence of WSDs is not completely explained by overall movement, and may therefore be related to other physiological artifacts such as respiratory variations (22).

### Identifiability of functional-connectivity networks

We next investigate how DiCER affects the correlation structure of rsfMRI time series, focusing particularly on whether DiCER over-corrects the data through iterative regression, thereby removing important neuronal contributions to rsfMRI signal. We conduct this investigation at two levels: (i) whole-brain functional connectivity (FC) matrices; and (ii) canonical functional connectivity networks identified using spatial independent component analysis (ICA). If DiCER is overly aggressive, we expect to see minimal (or distorted) FC structure, but if it successfully de-noises the data without being overly aggressive, we expect it to recover well-defined networks with comparable or enhanced sensitivity relative to other processing pipelines. We focus this analysis on the reduced (low motion) UCLA cohort. Results for the Beijing and Cambridge dataset are presented in the Supplementary Results (Figs S12, S13).

### Whole-brain networks

FC matrices are plotted for three different processing methods in Fig. 10. As previously reported (5, 15), processing using ICA-AROMA+2P retains WSDs that yield globally correlated FC structure, with minimal visible subnetwork structure (Figs 10A,D). This is in stark contrast to GMR (Fig. 10B), which yields a mean FC of zero by construction (18), resulting in a substantial proportion of negatively correlated sub-networks (Fig. 10E). DiCER recovers similar positive FC structure as GMR (Fig. 10C) but with a reduction in prominent anticorrelations (Fig. 10F). Thus, compared to GMR, DiCER recovers a similar network structure of positive functional connectivity, without inducing a large number of strong anticorrelations.

**Fig. 10.**
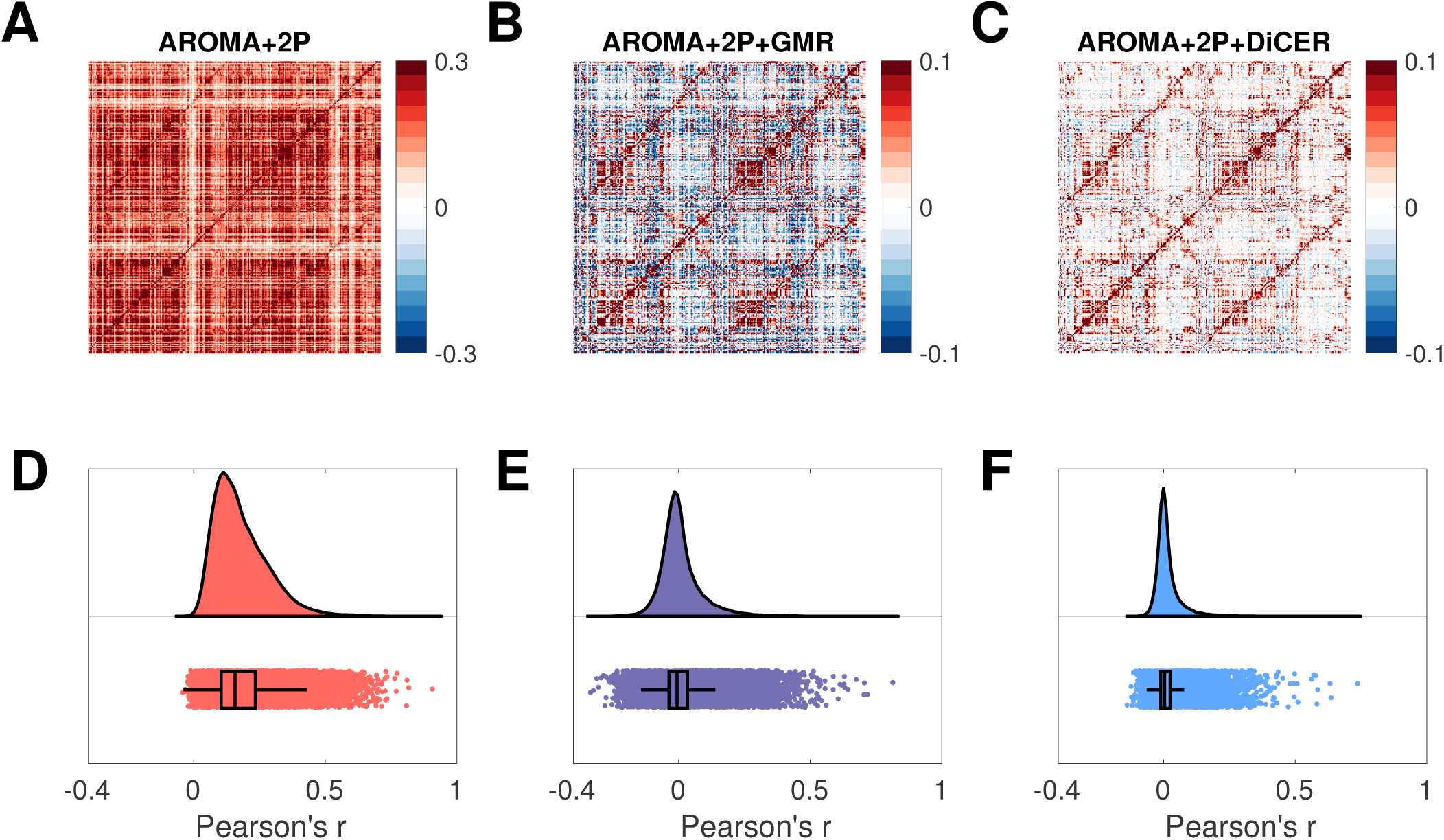
Functional connectivity (FC) depends strongly on fMRI preprocessing. We plot FC estimates for: **A** ICA-AROMA+2P, **B** ICA-AROMA+2P+GMR, and **C** ICA-AROMA+2P+DiCER. FC was estimated as the Pearson correlations between the mean signal in each pair of regions for a 333-ROI-per-hemisphere cortical parcellation (79). The group-level estimates shown here were computed by edge-wise averaging of the *z*-transformed FC matrices over all 121 UCLA control subjects. Nodes are ordered in the left hemisphere (upper half) followed by the right hemisphere (lower half). Note that the color bar range varies between plots: **A**: ±0.3, **B**–**C**: ±0.1. We also plot the distributions of Pearson’s correlation coefficients, *r* (after Z-transformation), for: **D** ICA-AROMA+2P, **E** ICA-AROMA+2P+GMR, and **F** ICA-AROMA+2P+DiCER, where the distributions are shown as smoothed kernel density estimates at top and boxplots at bottom, with medians and interquartile ranges annotated (80).

### Mapping canonical resting state networks with spatial ICA (sICA)

We next used sICA within the reduced UCLA control cohort as implemented in FSL’s `MELODIC` (32) to identify canonical resting-state networks. Because this dataset comes from two different scanner versions, we restricted our analysis to a subset of 89 subjects (note: these are within the reduced UCLA dataset, i.e., they are ‘low-motion’ subjects). Using independent component analysis (ICA), FSL Melodic linearly decomposed group-level 4D data into spatially independent components (ICs). This results in a number of spatial components that are maximally independent, along with their corresponding time course and power spectrum. Dual regression was then run to estimate subject-specific time series and associated spatial maps for each IC (81). The first stage of dual regression regressed each subject’s 4D data against the group averaged spatial maps from FSL Melodic. For each subject, this step yields the same number of time courses as the number of group-level spatial maps. The time courses are variance normalised to allow the investigation of both shape and amplitude of a resting-state network (81). In the second stage, each subject’s 4D data were then regressed against the component-specific time courses to estimate subject-specific spatial maps, one per group-level IC. We then estimated the strength of group-level networks by running FSL `Randomise` (82) permutation test on the subject-specific *t*-test statistical spatial maps, using 5000 permutations (otherwise know as stage 3 in a dual regression). Spatial ICs were classified post-hoc either ‘signal’, ‘noise’, or ‘unknown’ by analyzing their spatial maps, time series, and power spectra (13).

Using a Bayesian method to automatically determine the number of components to extract in the UCLA cohort (83), we find 123 components following AROMA+2P, 125 components following AROMA+2P+GMR, and 89 components following AROMA+2P+DiCER (for the Beijing cohort, the numbers were 176, 193 and 173, respectively). Different preprocessing pipelines will affect the inherent dimensionality of the residual data in different ways, and this will affect the final results of Melodic sICA. Interpreting these variations can be difficult. One the one hand, the presence of many components explaining small amounts of variance in high-dimension data could be consistent with high levels of residual noise; on the other hand, uncorrected WSDs may cause large-scale correlations that reduce intrinsic dimensionality. Nonetheless, some useful insight can be gained from inspecting Table 2, which labels the first 10 ICs identified following each processing pipeline in descending order of percentage of variance explained. Only one of the top ten ICs is identified as ‘signal’ following AROMA+2P or AROMA+2P+GMR, whereas AROMA+2P+DiCER yields nine signal components in the top ten. The ICs classified as ‘signal’ correspond to classic resting-state networks in all three denoising pipelines (13) (see Figs 11A,C,E for an example of the default-mode network). These results indicate that noise dominates the residual signal following AROMA+2P or AROMA+2P+GMR, whereas the majority of variance following DiCER is accounted for by ‘signal’ ICs.

**Table 2.** The first ten MELODIC components predominantly comprise signal ICs corresponding to classic functional networks following DiCER. In each denoising stream, bold font indicates a spatial IC component that corresponds to a classic resting-state network (13). This table is a subset of all networks, but here we identify: left and right fronto-parietal network (FPN), Cingulo-opercular network (CON), the default mode network (DMN), the visual network, and the sensorimotor network.

| IC component | ICA-AROMA+2P | ICA-AROMA+2P +GMR | ICA-AROMA+2P+DiCER |
| --- | --- | --- | --- |
| 1 | noise | noise | **left FPN** |
| 2 | noise | noise | **CON** |
| 3 | noise | noise | **DMN** |
| 4 | noise | noise | **visual** |
| 5 | noise | noise | **left FPN** |
| 6 | noise | noise | **right FPN** |
| 7 | noise | noise | **visual** |
| 8 | **visual** | noise | noise |
| 9 | noise | **DMN** | **DMN (post)** |
| 10 | noise | noise | **sensorimotor** |

**Fig. 11.**
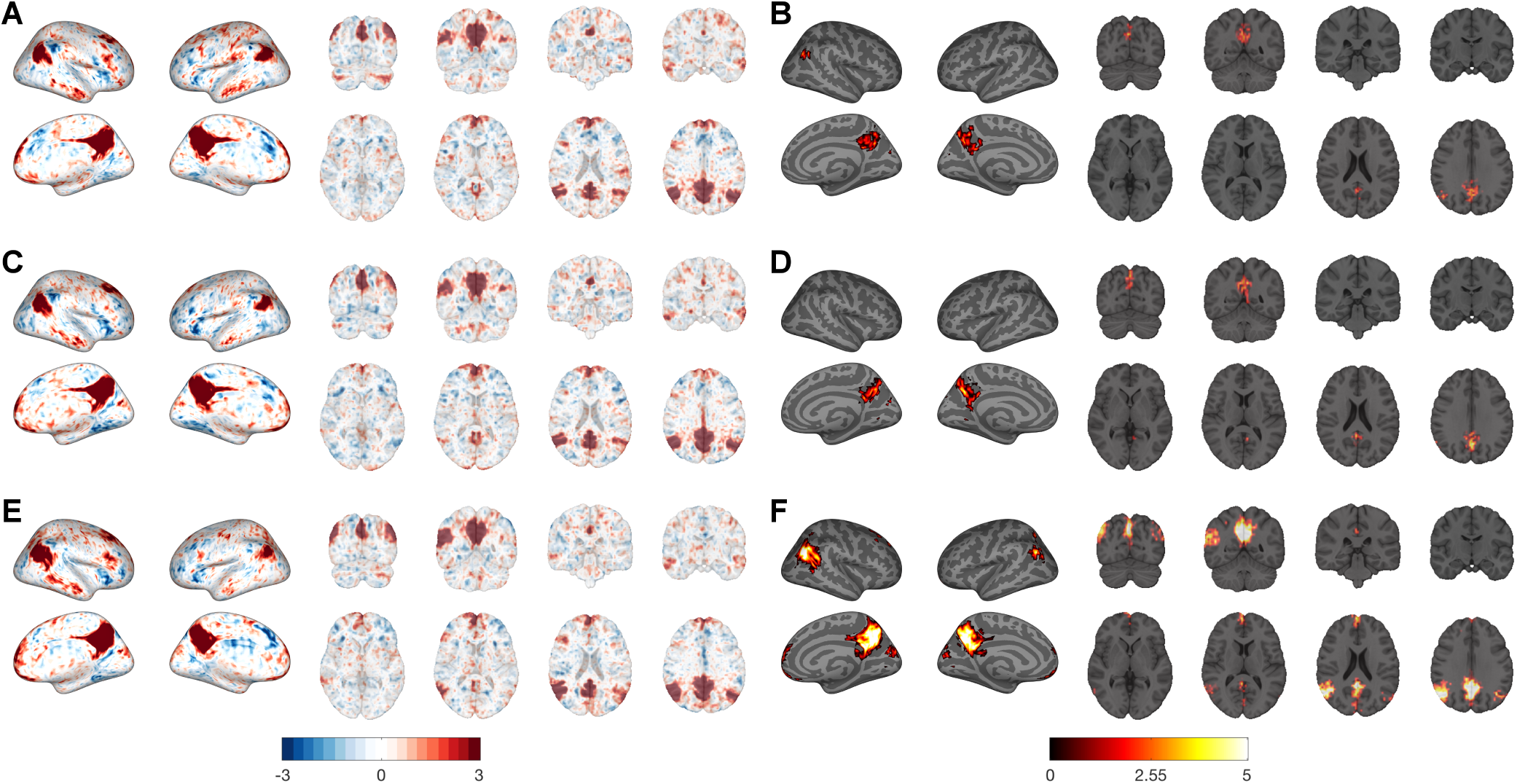
Identifiability of default-mode network (DMN) using Melodic ICA following different preprocessing pipelines. The group-level independent component maps, with the corresponding activation map for AROMA+2P, AROMA+2P+GMR, and AROMA+2P+DiCER are shown in the pairs **A**,**B, C**,**D**, and **E**,**F** and respectively. The group-level activation maps are calculated from dual-regression. The *t*-statistic maps have been thresholded at p<.05, FWE-corrected via Randomize.

Comparison of IC spatial maps obtained across pipelines is complicated by differences in the intrinsic dimensionality of the denoised data. With low-dimensionality data, networks typically load onto fewer components, resulting in some classic networks merging into single components. When the dimensionality of the data is sufficiently high, classic networks can split into two or more components. Here, we compared specific ‘signal’ ICs that clearly corresponded to classic resting-state networks, and which showed a similar spatial topography across the three pipelines. For the UCLA cohort, we identified five such networks. Spatial maps for one canonical IC, the default mode network (DMN), are plotted following AROMA+2P (Fig. 11B), AROMA+2P+GMR (Fig. 11D), and AROMA+2P+DiCER (Fig. 11F). Maps and corresponding results for the other ICs are shown in Figs S13–S20 (the results for six comparable ICs identified in the Beijing cohort are show in Figs S21–S32). Figure 2 shows second-level maps of voxels showing statistically significant contributions to the IC (*p* < 0.05 corrected as described above) across the entire group. The DMN component’s spatial structure is broadly similar across the three pipelines (also indicated by the underlying similarity of the ICs in Figs 11A,C,E), but the spatial clusters comprising the network are generally larger in extent following DiCER. It is also evident that the *t*-statistics are higher in DiCER, an effect more clearly visualized in Fig 12 where we how the distribution of *t*-statistics for the unthresholded network maps identified following each processing stream. The maximum *t*-statistics are similar for AROMA+2P (5.95) and AROMA+2P+GMR (5.65). In contrast, the distribution for AROMA+2P+DiCER shows an extended tail to a maximum of 10.35, which is nearly double the maxima of the other two pipelines. Similar results were obtained for the other canonical resting-state networks identified as ‘signal’ ICs in our analysis for both the UCLA and Beijing data. The differences between pipelines were smaller in the Beijing data, perhaps due to the low levels of motion and noise in this cohort. Nonetheless, in nearly all cases, DiCER performed as well as, or better, than GMR (see Supplementary Text 2 for results for other ICs in the UCLA and Beijing datasets). Together, these results indicate that DiCER results in stronger identification of classic functional-connectivity networks, likely due to its efficacy in reducing inter-individual variance in residual noise (shown in Fig. 9).

**Fig. 12.**
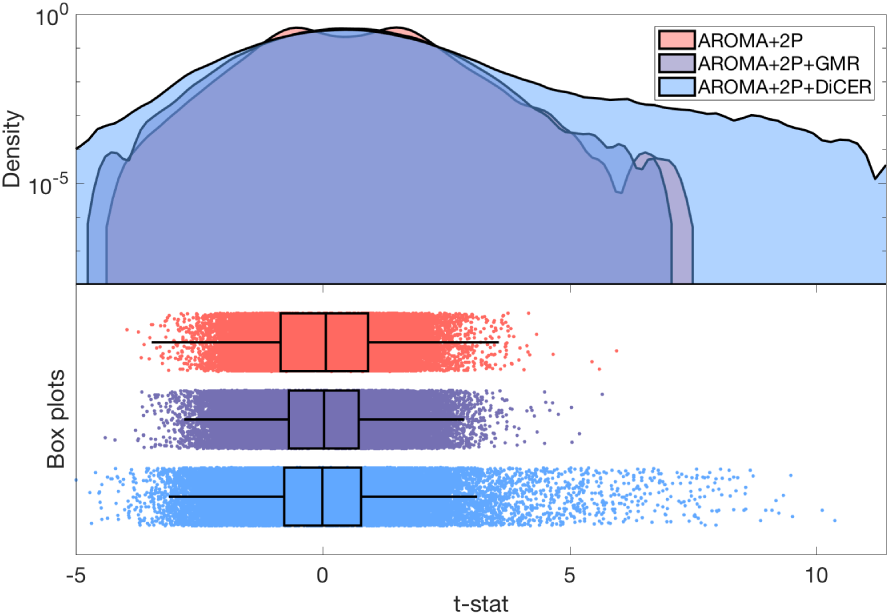
Group-level *t*-statistic distributions for the DMN, identified through sICA, following different denoising pipelines. Note, the logarithmic vertical axis for the smoothed kernel estimate (top panel) to emphasize the tails of this distribution.

## Discussion

The main conclusions of our analyses are summarized in the following.

1. The way in which a carpet plot is ordered can have a major impact on how one understands the spatiotemporal structure of fMRI data.
2. Data-driven reordering of voxels (or regions) in a carpet plot (e.g., CO) reveals diverse WSDs that are apparent across time within an individual and betweeen different individuals. These WSDs can be classified as either monophasic or biphasic, and they vary in their intensity and spatial extent.
3. GMR is only effective in removing some of these WSDs; namely, monophasic WSDs with a broad anatomical distribution. The limitations of GMR are only clearly apparent when inspecting CO (or GSO) carpet plots.
4. We introduce a new fMRI data-correction method, DiCER, which successfully removes diverse WSDs without imposing a specific structure on the corrected data (e.g., a zero-centered distribution of pairwise correlations (18)).
5. DiCER improves commonly used quality-control benchmarks (e.g., QC–FC correlations), reduces inter-individual variability in global correlation structure, and results in superior identification of canonical resting-state networks using popular methods such as sICA, with an approximate two-fold increase of statistical sensitivity for some networks.

### Carpet-plot ordering and the diversity of WSDs

Our findings underscore the critical benefits of appropriately ordering carpet plots when investigating the spatiotemporal structure of fMRI datasets. Conventional voxel orderings (labeled as ‘RO’ here) often fail to reveal the full range and structure of WSDs. Reordered carpet plots allow WSDs to be classified as monophasic or biphasic. Individuals vary in their expression of monophasic and biphasic deflections: some exhibit predominantly monophasic WSDs, some show mainly biphasic WSDs, and some show a mixture. Whether these distinct classes of WSDs arise from different causes is unknown. Both monophasic and biphasic WSDs can be timelocked to spikes in FD traces, but can also arise independently of large head movements. It is possible that these additional WSDs may be caused by respiration (22); examining their relation to both respiratory and other physiological traces is an important avenue of further work.

### On the efficacy of GMR (and GSR)

Carpet-plot ordering is critical when evaluating denoising efficacy. As shown in Fig. 1, the default ordering (RO) suggests that GMR is quite effective in removing WSDs. However, when the plots are reordered according to either GSO or CO, it is apparent that GMR leaves considerable residual structure. Biphasic WSDs are particularly immune to the effects of GMR. This result can be understood by considering the extreme case in which half the brain shows a positive signal deflection and the other half shows a negative signal deflection of equal magnitude: the global mean at such a time point will be zero and will thus contain no information about the biphasic deflection (and nothing will be corrected). GMR is effective in cases where there is an anatomically widespread, monophasic WSD, but our results suggest that approximately one third of individuals exhibit prominent biphasic WSDs, with even more exhibiting a mix of monophasic and biphasic WSDs. As a result, the efficacy of GMR (or GSR) in removing WSDs varies considerably across individuals, as shown in Fig. 9. This variable efficacy leaves unwanted, noise-related inter-individual variance in the data, limiting sensitivity for identifying functional networks (e.g., Fig. 11 and Fig. 12).

A key step in DiCER involves computing the adjusted mean, which attempts to align all voxel time courses by introducing a sign flip to voxels with anticorrelated time courses. This sign flip aims to address the problem posed by anticorrelated signals within a cluster, ensuring that, regardless of their direction, common fluctuations contribute additively to the mean rather than cancelling out. This step enables more effective correction of biphasic WSDs.

### On the efficacy and implementation of DiCER

DiCER is designed to capture diverse WSDs with a broad anatomical distribution through an iterative estimation and removal approach. As shown in Fig. 4, our approach has considerable flexibility in capturing signals with a range of temporal characteristics, including those with low-frequency trends and bursty signal deflections. This flexibility results in successful flattening of carpet plots, even when visualized using GSO or CO: it dramatically reduces inter-individual variability in global correlation structure, as quantified using VE1 (Fig. 9) and improves quality metrics (QC–FC and reduces the ‘negative’ dip in the distance-dependence). Collectively these results indicate that AROMA+2P+DiCER is more effective than either AROMA+2P or AROMA+2P+GMR in removing WSDs and reducing the motion-dependence of FC estimates. Our implementation of DiCER requires the setting of three key parameters (`dbscan`: *N*_points_, ϵ, and the maximum number of iterations, *k*_max_), which we set in order to estimate diffuse, common signals in rsfMRI data. We also enforced a minimum cluster size (at least 10% of the total number of voxels need to be in the core of a cluster for it to be considered further). These parameters were set manually through empirical testing on the UCLA, Beijing, and Cambridge datasets. The degree to which these settings generalize to other datasets remains unclear, as does the dependence of DiCER performance on the preprocessing steps that precede it.

Future work could attempt to explicitly fit these parameters to data in a way that aims to strike a balance between regimes of under-correcting (stringent parameters) and over-correcting (lenient parameters), and further test their generalizability to diverse fMRI datasets. There are several addtional, straight-forward directions in which the method could be further developed. For example, we included all voxels in the cluster in estimating the adjusted mean, but the clustering approach used in DiCER gives additional information about the contribution of different voxels to the identified WSD, labeling voxels as either core or reachable, and providing a measure of ‘distance’ from the cluster center. This information could be used to reweight voxels to more accurately compute a representative WSD signal for an identified `dbscan` cluster.

A particular strength of DiCER is that it need not identify regressors when it finds no evidence for WSDs (i.e., when there are no dense clusters of voxels in the defined similarity space). In principle, this means that we apply no correction to individuals who do not show prominent WSDs. In practice, with the settings used here, the number of regressors identified for any given individual varied between 1 and 5 across the three datasets that we studied. Tuning the settings for cluster definition can change the number of regressors identified, thus providing investigators with measurable control to determine the degree to which they denoise their data. Indeed, alhtough our parameter settings generally worked well and out-performed GMR, in some cases residual, small-scale WSDs were apparent (e.g., sub-10934 in Fig S6 and sub-10321,sub-10460 in the report https://bmhlab.github.io/DiCER_results/). These could be removed through further iterations of DiCER. The flexibility of our approach contrasts with GMR/GSR, where a single regressor is applied to every dataset. A remaining challenge is to develop a principled, data-driven method for determining optimal algorithm settings.

### Denosing fMRI data: how far is too far?

A challenge for evaluating fMRI denoising methods is that we lack a gold standard for distinguishing signal from noise. It is therefore difficult to determine whether a proposed denoising signal is going too far by removing real neural signal. This is a salient concern for a method such as DiCER, which is iterative and results in a flattened carpet plot that visually lacks obvious spatiotemporal structures (most clearly visualized in CO carpet plots where the voxel ordering is determined based on the cleaned data, shown at https://bmhlab.github.io/DiCER_results/). Indeed,our approach assumes that signals driving coherent fluctuations in large sets of voxels are artifactual. This assumption is justified by evidence tying WSDs to artifacts related to scanner issues, motion, and respiration (22, 25), but there is some debate about whether all WSDs should be treated as noise (19, 20). Techniques such as tICA may be more effective in distinguishing WSDs related to noise from those related to neuronal dynamics (46), but is presently only applicable to very-high resolution data. Our analyses suggest that, in the worst case scenario, DiCER is not more aggressive than GMR in removing neural signal. In fact, our sICA analysis suggests that DiCER greatly improves the identifiability of canonical resting-state networks, often leading to two-fold gains in statistical sensitivity. It is also more effective at denoising, given that 9/10 of the top sICA components were identified as ‘signal’ following AROMA+2P+DiCER, cf. Table 2), whereas the bulk of the fMRI variance following AROMA+2P and AROMA+2P+GMR is accounted for by noise (9/10 of the leading sICA components were labeled ‘noise’). When taken with the QC-FC findings, these results suggest that, rather than removing too much signal, DiCER is successfully denoising the data and improving the detection of brain networks. Complimentary strategies, such as considering how DiCER influences task-evoked activations or brain-behaviour correlations (43) (although see also Siegel et al. (84)), may prove useful in further validating our approach.

Lastly, we note that in the case where the data is relatively ‘clean’ (with respect to WSDs and motion), such as in the Beijing cohort, IC networks are still discovered with ICA with comparable or improved sensitivity following DiCER compared to AROMA+2P or AROMA+2P+GMR. Indeed, the top 10 ICs were designated ‘signal’ following DiCER, compared to 3/10 for the other two pipelines (see Supplementary Text 2). These results provide further confidence that DiCER has not over-corrected the data relative to GMR.

## Conclusions

Removing WSDs from rsfMRI data is a challenge for many denoising algorithms. GSR is a controversial procedure that is thought to successfully remove WSDs at the expense of imposing a specific structure on the residual data and possibly distorting connectivity estimates (16, 18). Here we show that WSDs have a more complex structure than previously thought, that GSR is effective only in removing some of these WSDs, and that typical methods for visualizing the data (i.e., RO carpet plots) mask WSDs and their diversity. We introduce DiCER, a new iterative approach that successfully removes diverse kinds of WSDs, reduces correlations between functional connectivity and head motion, reduces inter-subject variability in global correlation structure, and improves statistical sensitivity for identifying canonical functional-connectivity networks.

## ACKNOWLEDGEMENTS

AF was supported by the Australian Research Council (ID: FT130100589), National Health and Medical Research Council (NHMRC; ID: 1104580), and the Sylvia and Charles Viertel Charitable Foundation. BDF was supported by an NHMRC Fellowship (1089718).

