## Supplementary Text 1 for "Identifying and removing widespread signal deflections from fMRI data: Rethinking the global signal regression problem"

### Supplementary Text 1: ‘Identifying and removing widespread signal deflections from fMRI data: Rethinking the global signal regression problem.’

**Kevin M. Aquino**[1,*], **Ben D. Fulcher**[2], **Linden Parkes**[1], **Kristina Sabaroedin**[1], **and Alex Fornito**[1]

[1]The Turner Institute for Brain and Mental Health, School of Psychological Sciences, and Monash Biomedical Imaging, Monash University, Victoria, Australia.
[2]School of Physics, The University of Sydney, NSW 2006, Australia.

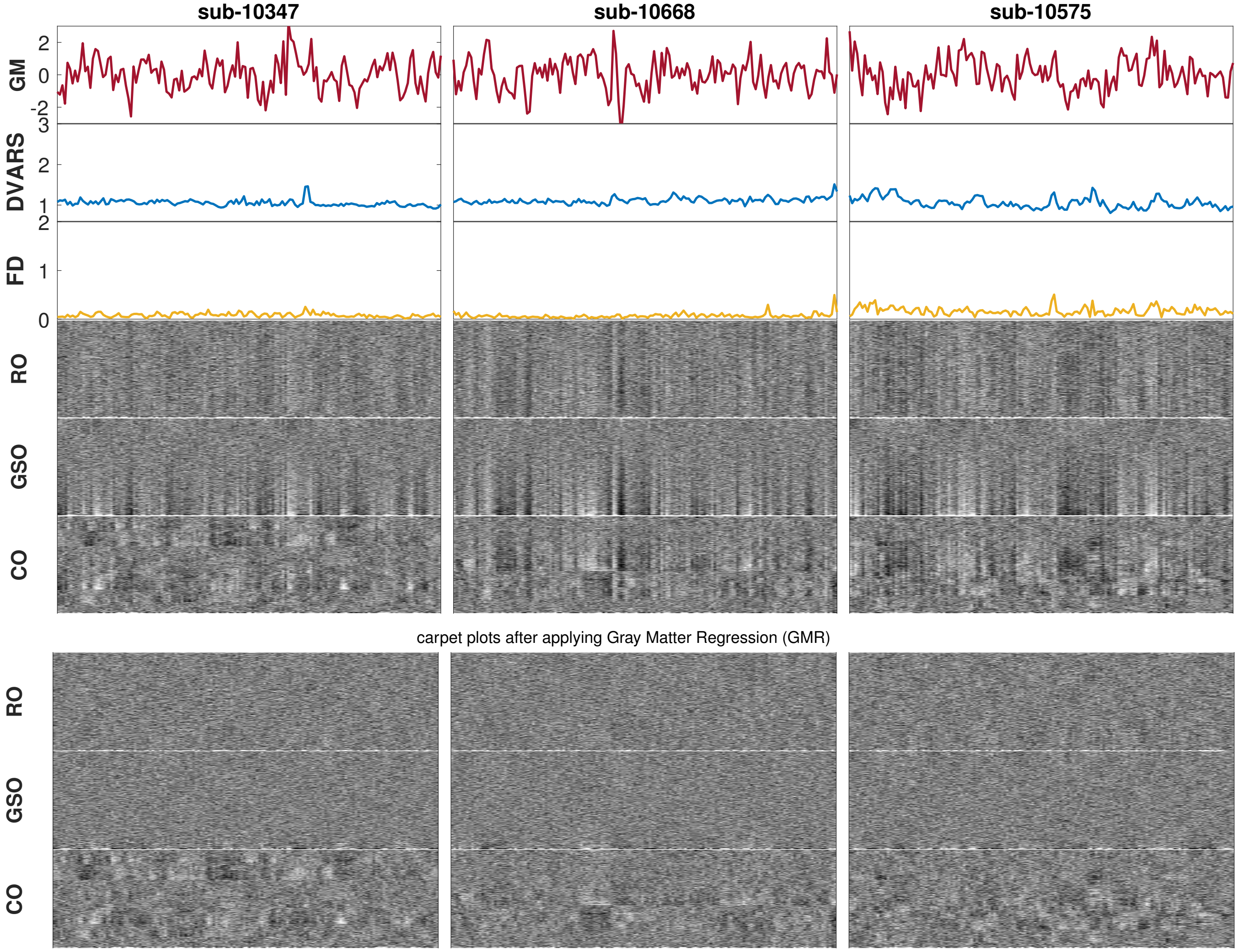


**Fig. S1. Exemplar Category I subjects (weak global signal)** Each column displays a different subject The first three rows show: (i) the mean GM signal, (ii) the temporal Derivative of root mean square VARiance over voxelS (DVARS)(1), and (iii) framewise displacement (FD). DVARS and GS are calculated after basic fmriprep preprocessing (before AROMA). The next three rows show carpet plots of the same data plotted using different voxel orderings: (iv) random order (RO) carpet plots; (v) global signal order (GSO) carpet plots; and (vi) cluster order (CO) carpet plots. All heatmaps use the same color scale and present identical data for each individual. The last three rows show the carpet plots after applying GMR (vii),(viii) and (ix) under RO,GSO, and CO respectively.

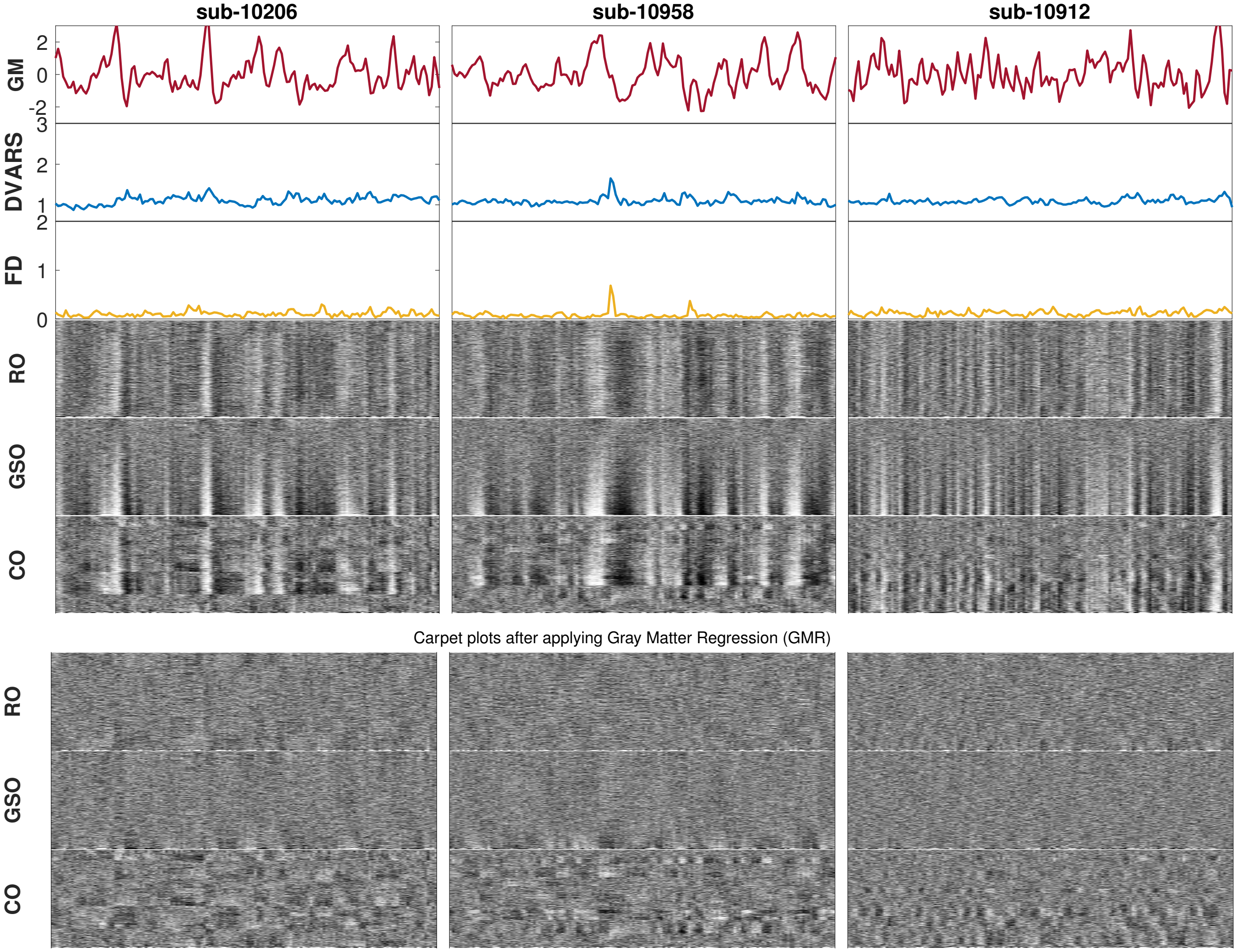


**Fig. S2. Category II subjects**: Each column displays a different subject. The first three rows show: (i) the mean GM signal, (ii) the temporal Derivative of root mean square VARiance over voxelS (DVARS)(1), and (iii) framewise displacement (FD). DVARS and GS are calculated after basic fmriprep preprocessing (before AROMA). The next three rows show carpet plots of the same data plotted using different voxel orderings: (iv) random order (RO) carpet plots; (v) global signal order (GSO) carpet plots; and (vi) cluster order (CO) carpet plots. All heatmaps use the same color scale and present identical data for each individual. The last three rows show the carpet plots after applying GMR (vii),(viii) and (ix) under RO,GSO, and CO respectively.

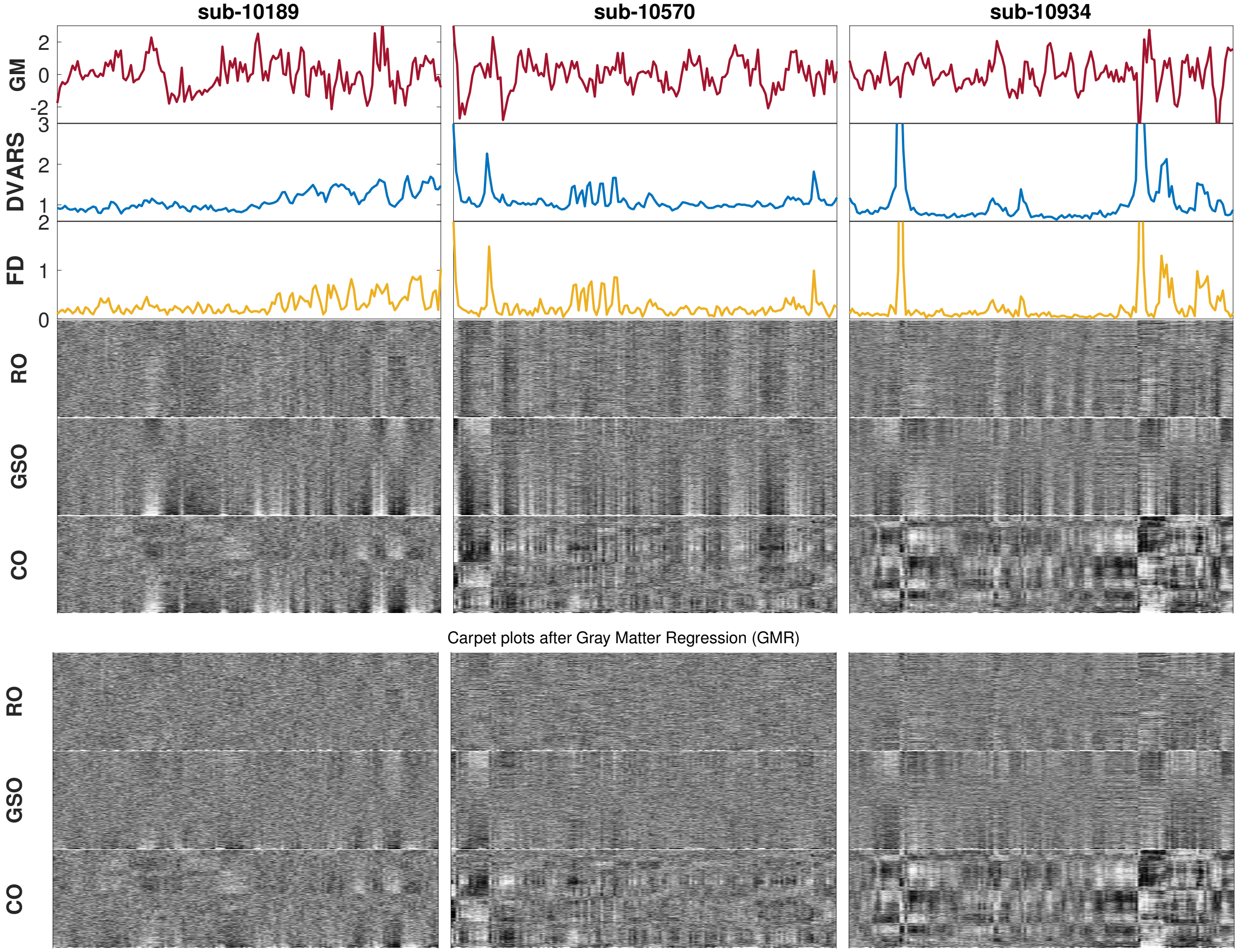


**Fig. S3. Category III Subjects** exhibit prominent biphasic deflections. Each column displays a different subject. The first three rows show: (i) the mean GM signal, (ii) the temporal Derivative of root mean square VARiance over voxelS (DVARS)(1), and (iii) framewise displacement (FD). DVARS and GS are calculated after basic fmriprep preprocessing (before AROMA). The next three rows show carpet plots of the same data plotted using different voxel orderings: (iv) random order (RO) carpet plots; (v) global signal order (GSO) carpet plots; and (vi) cluster order (CO) carpet plots. All heatmaps use the same color scale and present identical data for each individual. The last three rows show the carpet plots after applying GMR (vii),(viii) and (ix) under RO,GSO, and CO respectively.

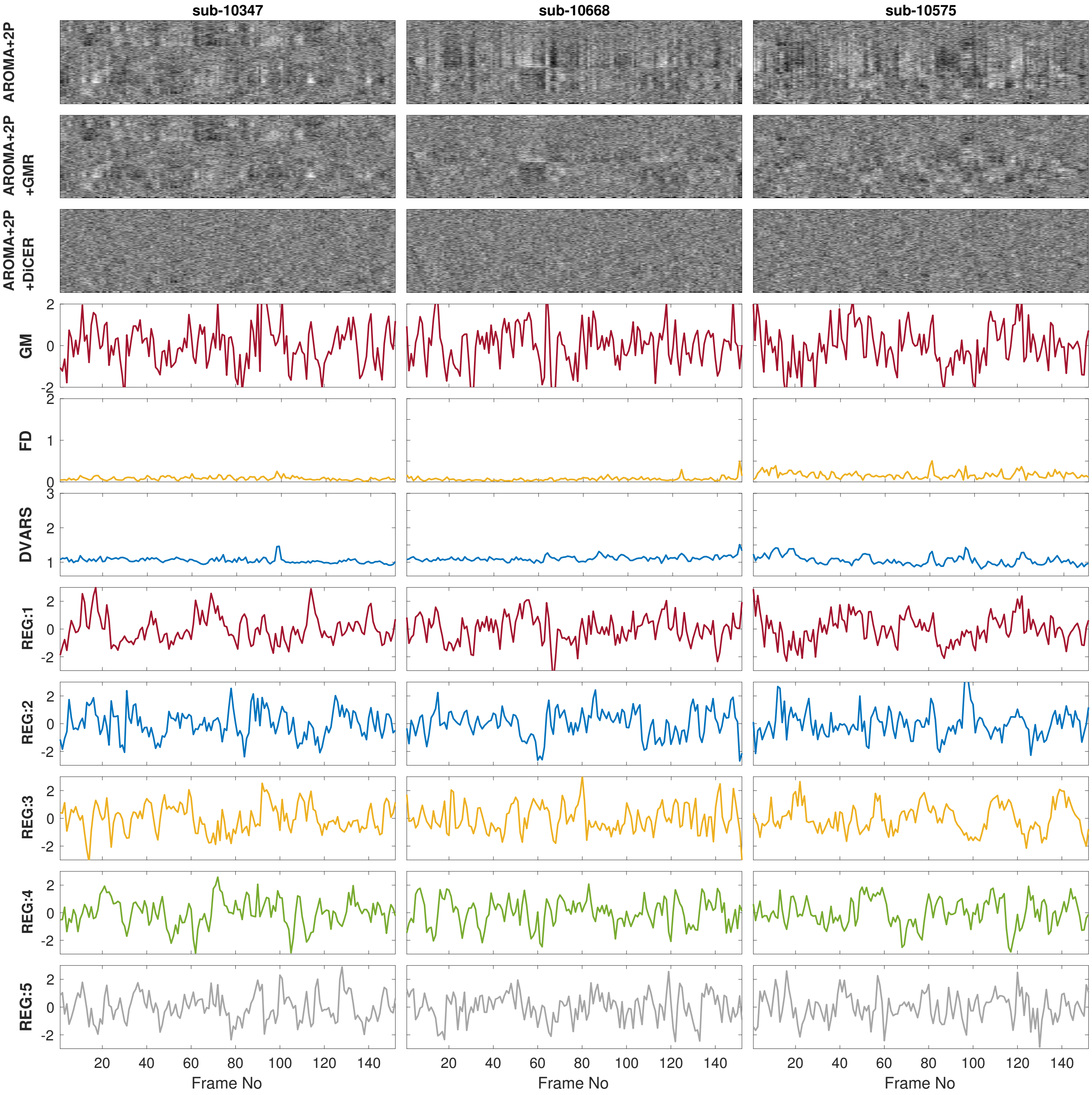


**Fig. S4. DiCER applied to three Category I subjects** The top three rows depict results obtained using three noise-correction schemes: (i) ICA-AROMA+2P, (ii) ICA-AROMA+2P+GMR, and (iii) ICA- AROMA+2P+DiCER, plotted as CO carpet plots. The fourth (iv) row shows the mean grey matter signal, the fifth (v) row shows FD and and the sixth row (vi) shows DVARS. The rows (vii)–(xi) are the regressors estimated by DiCER.

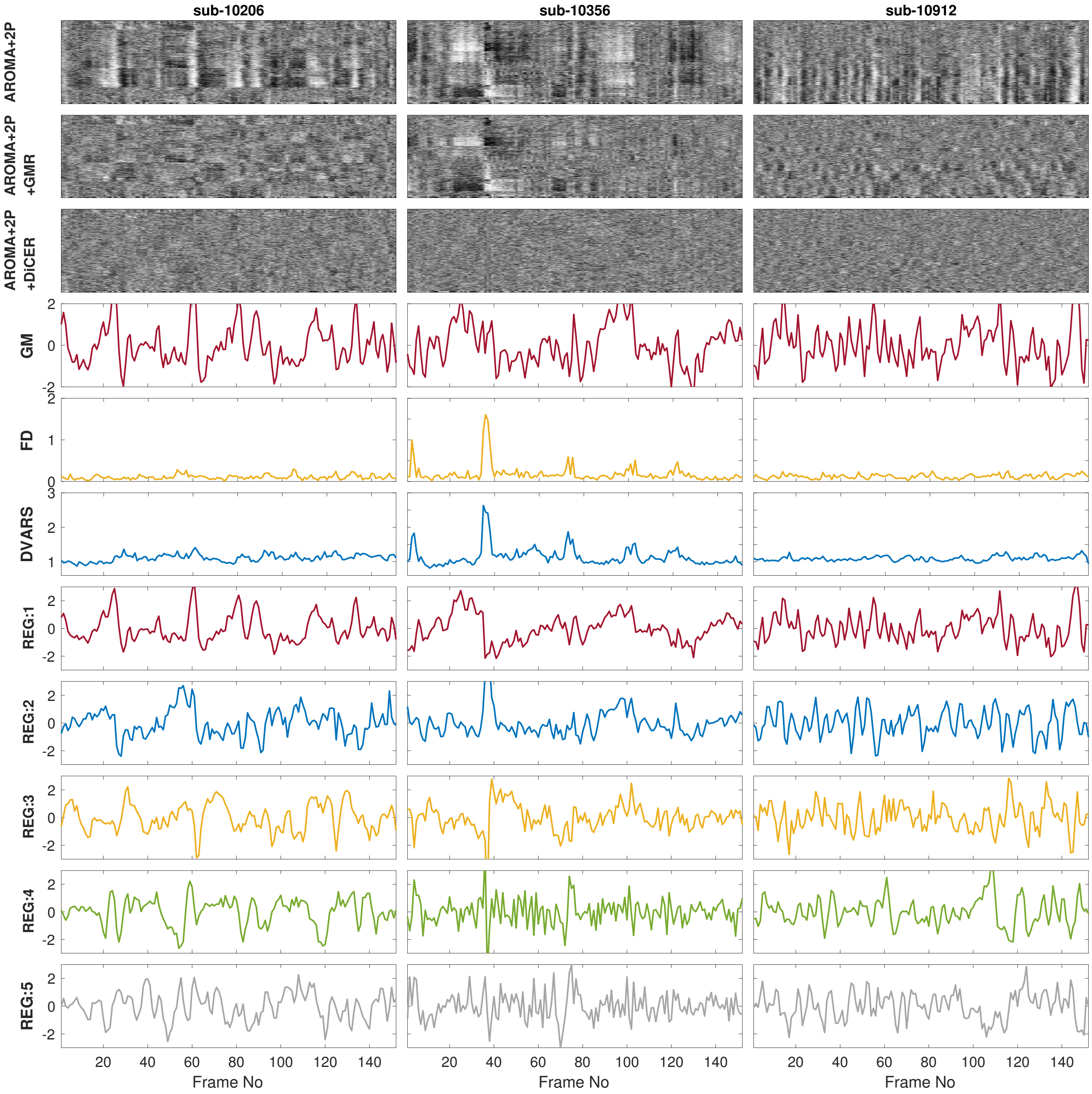


**Fig. S5. DiCER applied to Category II subjects**: The top three rows depict results obtained using three noise-correction schemes: (i) ICA-AROMA+2P, (ii) ICA-AROMA+2P+GMR, and (iii) ICA- AROMA+2P+DiCER, plotted as CO carpet plots. The fourth (iv) row shows the mean grey matter signal, the fifth (v) row shows FD and and the sixth row (vi) shows DVARS. The rows (vii)–(xi) are the regressors estimated by DiCER.

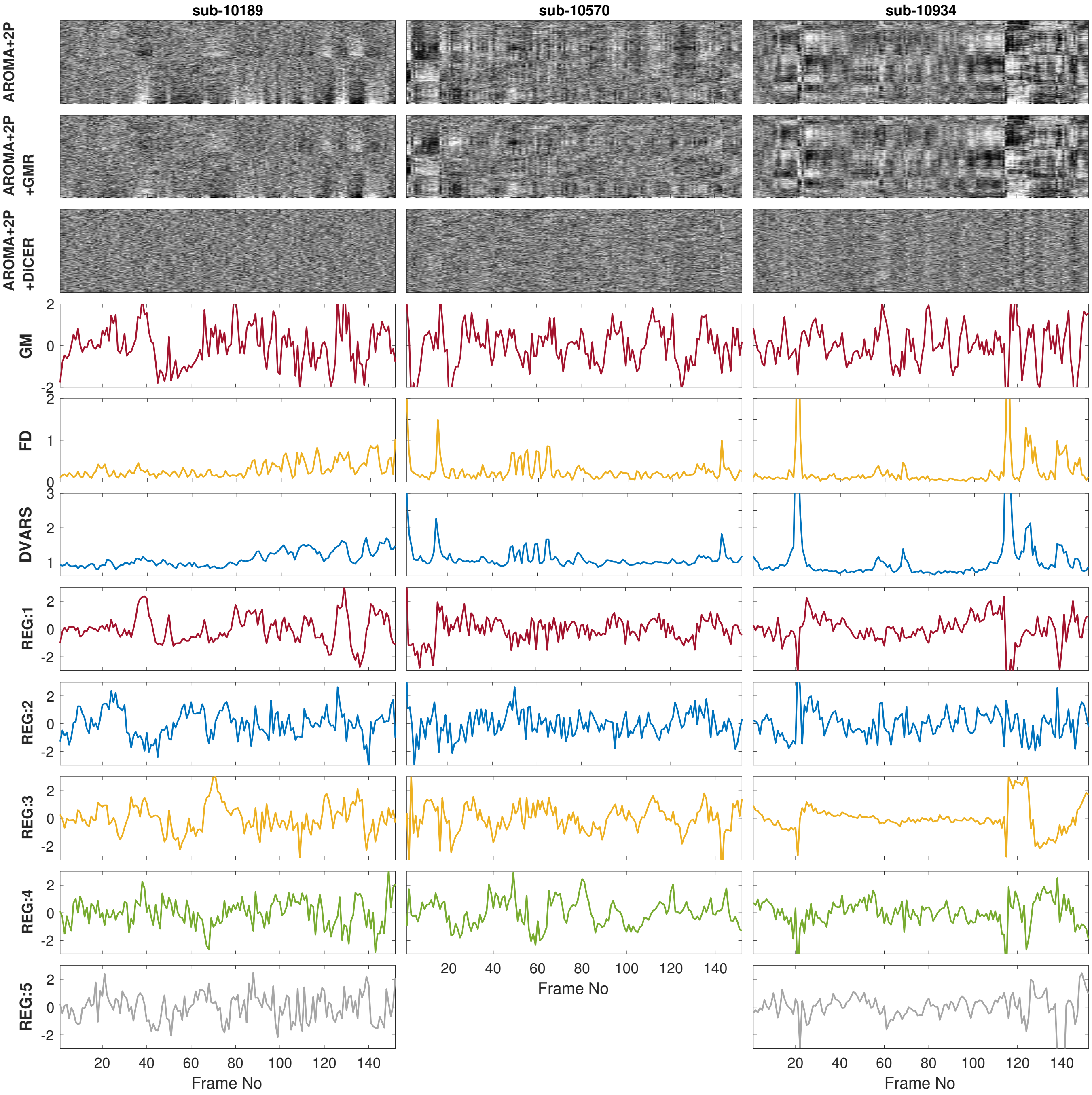


**Fig. S6. DiCER applied to Category III subjects** The top three rows depict results obtained using three noise-correction schemes: (i) ICA-AROMA+2P, (ii) ICA-AROMA+2P+GMR, and (iii) ICA- AROMA+2P+DiCER, plotted as CO carpet plots. The fourth (iv) row shows the mean grey matter signal, the fifth (v) row shows FD and and the sixth row (vi) shows DVARS. The rows (vii)–(xi) are the regressors estimated by DiCER.

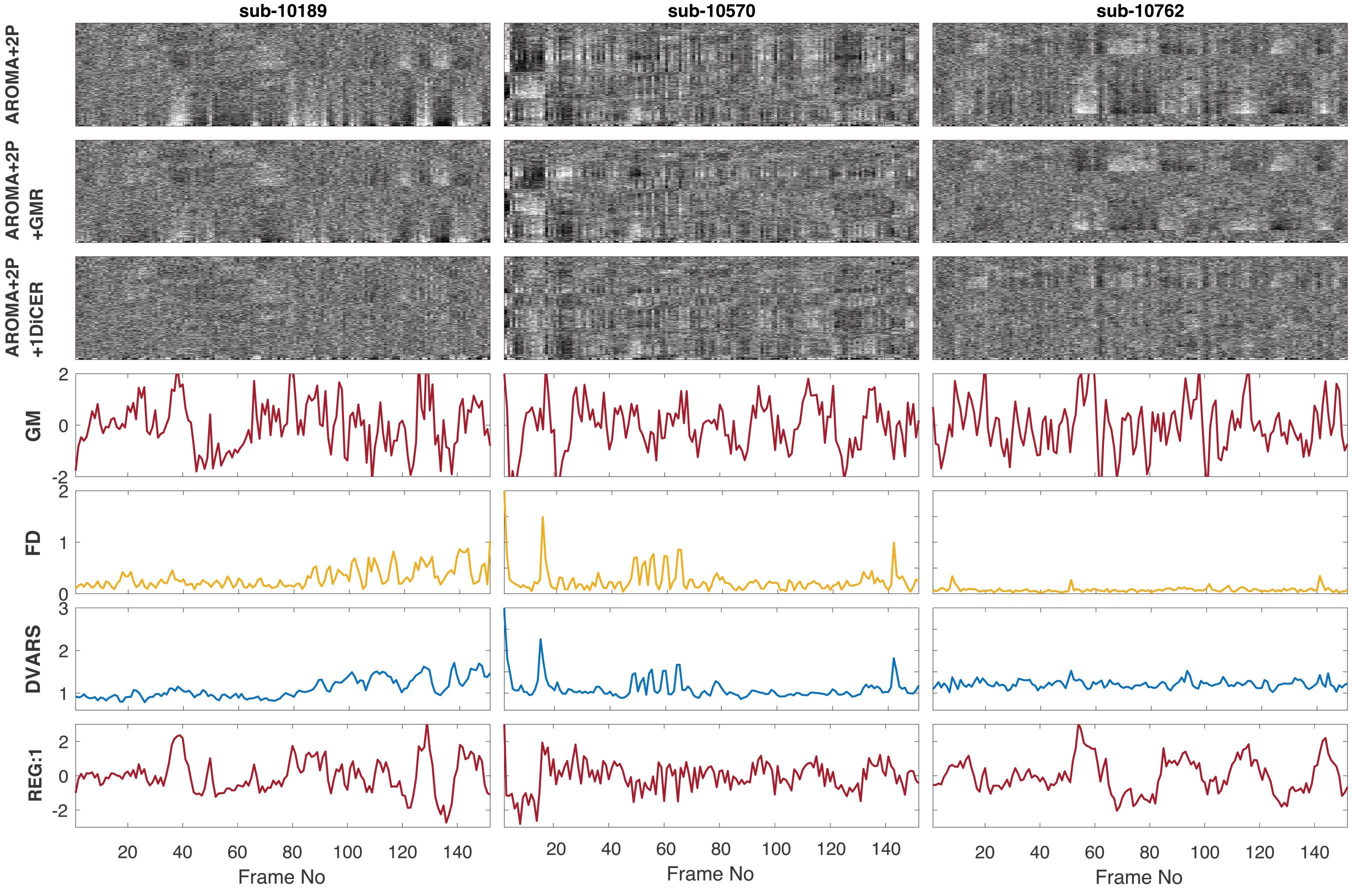


**Fig. S7. DiCER with 1 regressor applied to three subjects** The top three rows depict results obtained using three noise-correction schemes: (i) ICA-AROMA+2P, (ii) ICA-AROMA+2P+GMR, and (iii) ICA- AROMA+2P+1DiCER, plotted as CO carpet plots. The 1DiCER pipeline is DiCER with only 1 iteration (hence only one regressor). The fourth (iv) row shows the mean grey matter signal, the fifth (v) row shows FD and and the sixth row (vi) shows DVARS. The row (vii) is the first regressor estimated by DiCER.

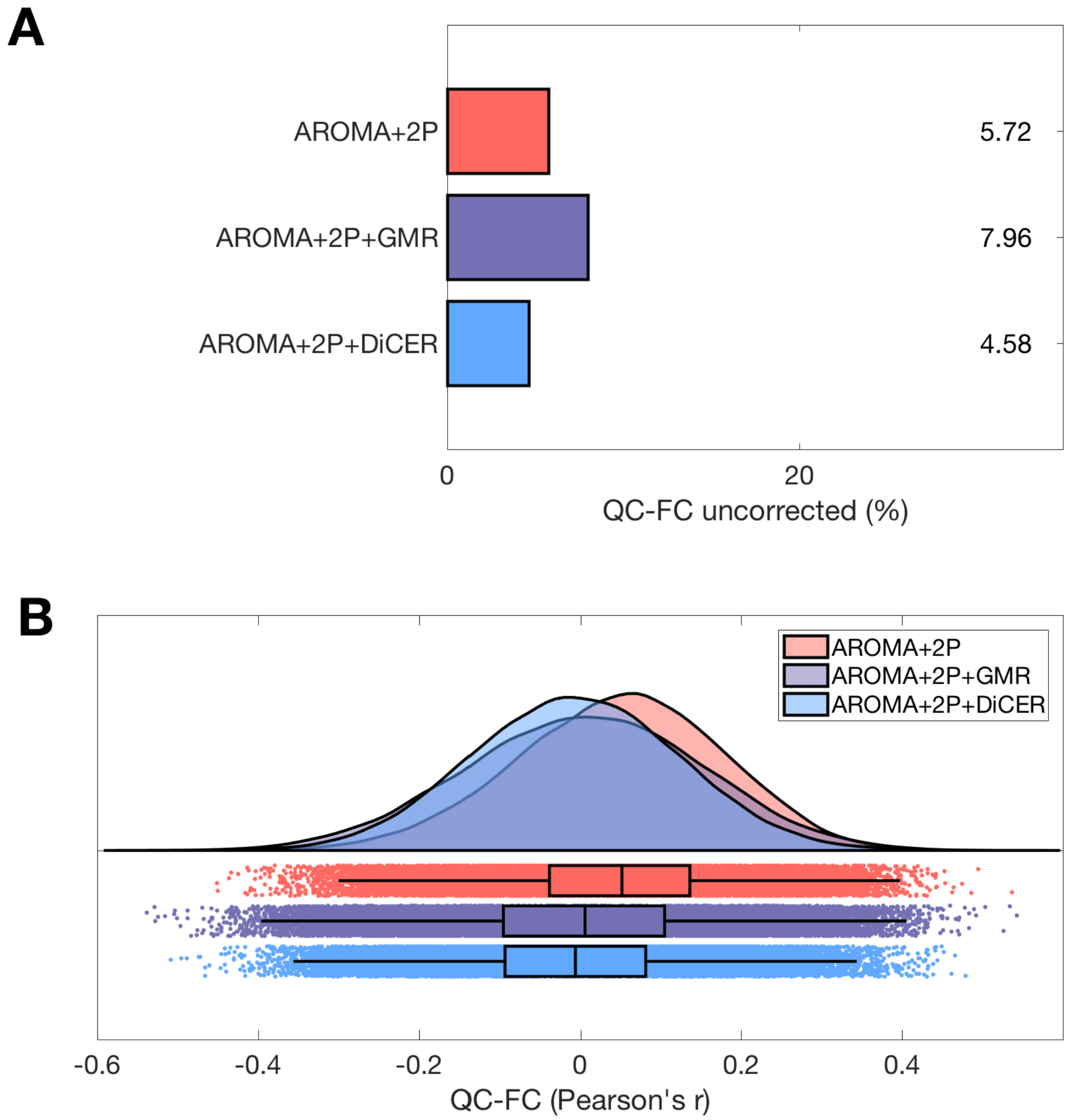


**Fig. S8. DiCER reduces the correlation between motion (mean FD) and functional connectivity (FC).** We compare ICA-AROMA+2P (red), ICA-AROMA+2P+GMR (purple), and ICA-AROMA+2P+DiCER (blue) on the reduced (low-motion) **A**, **B** Cambridge cohort. **A**, The proportion of FC values that are correlated to mean FD at a threshold of $p < 0.05$, uncorrected. **B**, The distributions of QC–FC correlations (Pearson's $r$) across all edges shown as smoothed kernel density estimates at top and boxplots at bottom, with medians and interquartile ranges annotated(2).

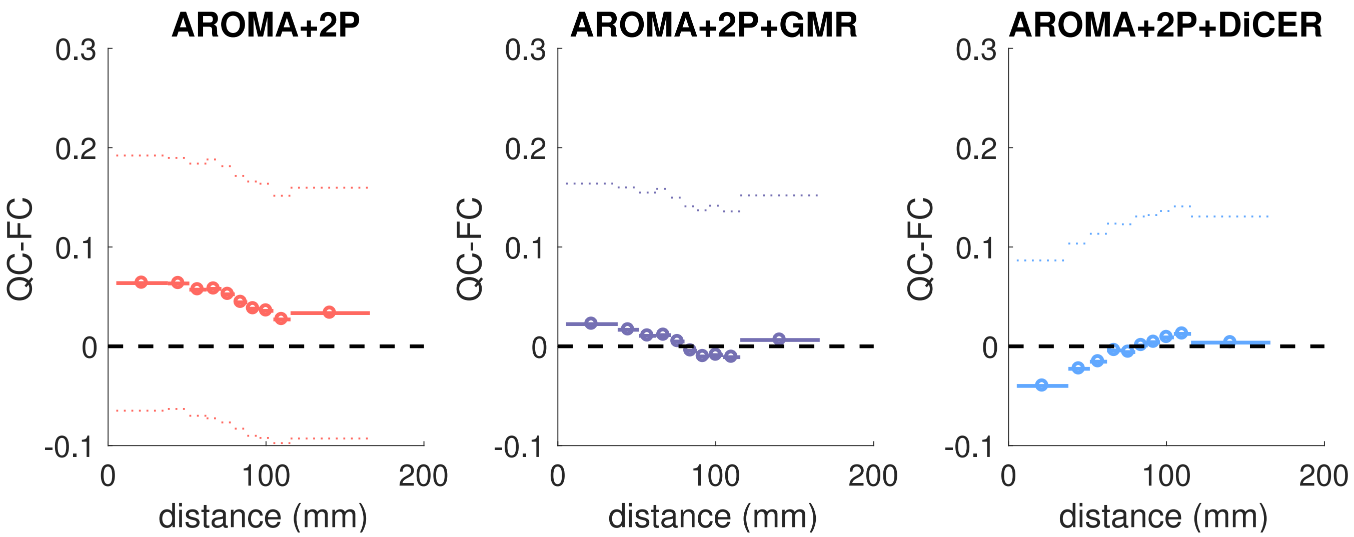


**Fig. S9. DiCER reduces the distance-dependence of QC–FC.** QC–FC is plotted across ten equiprobable distance bins for the Cambridge reduced low motion cohorts, shown as mean (circle and solid horizontal lines) ± standard deviation (dashed horizontal lines), for: ICA-AROMA+2P, ICA-AROMA+2P+GMR, ICA-AROMA+2P+DiCER.

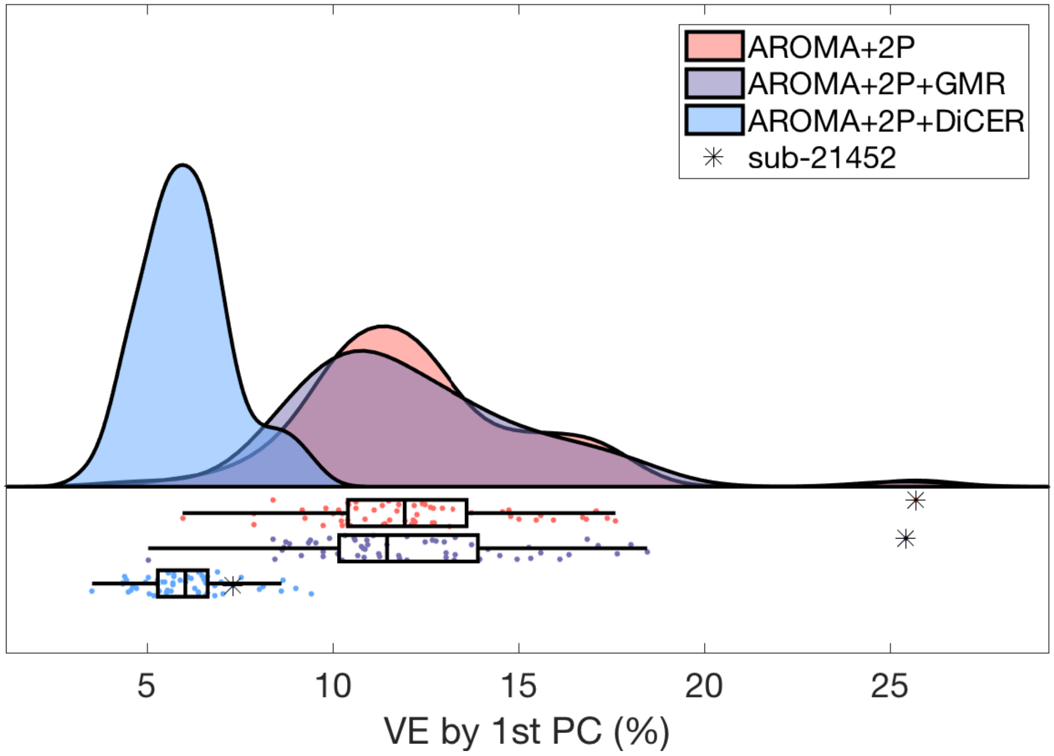


**Fig. S10.** The variance explained (% of total variance) by the first principle component of the carpet plots. The distributions of VE1 are shown as smoothed kernel density estimates at top and boxplots at bottom, with medians and interquartile ranges annotated (2).

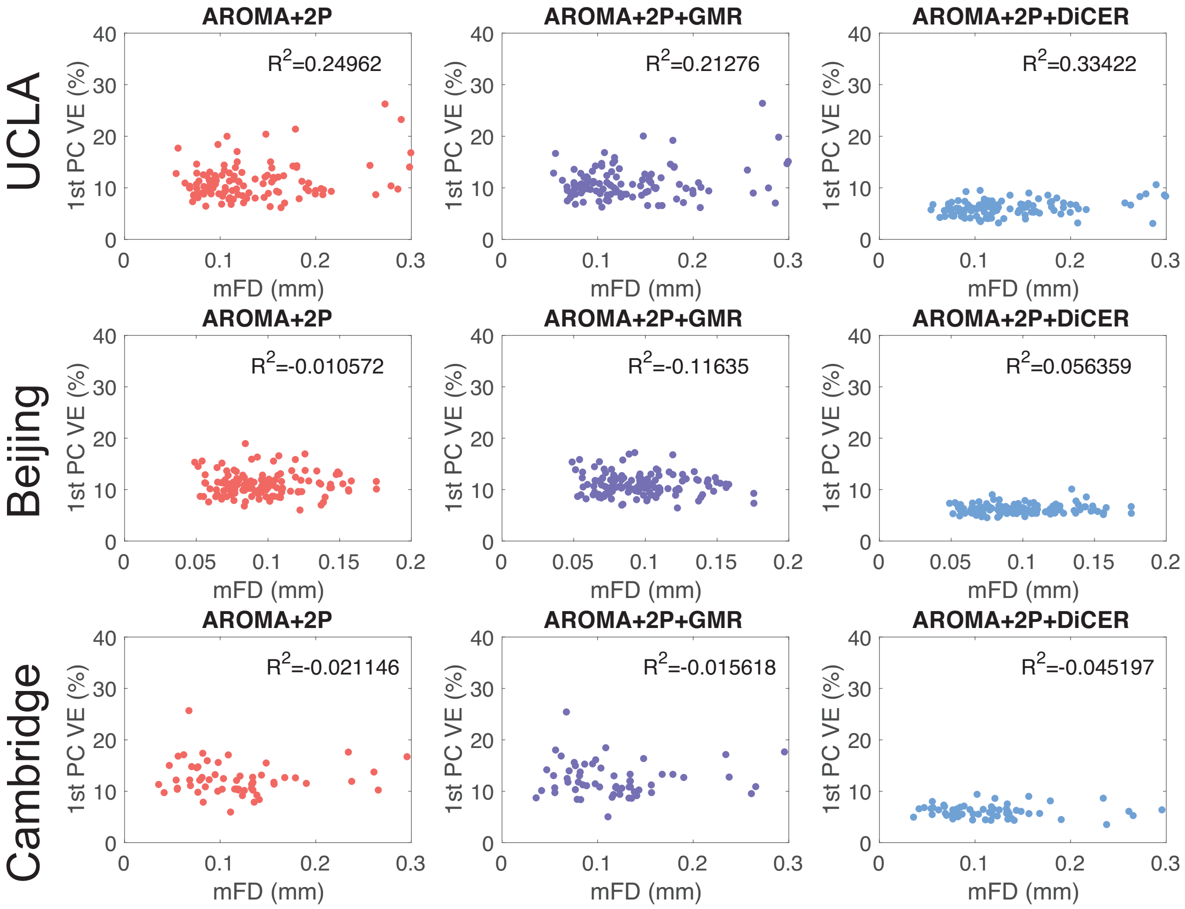


**Fig. S11.** Mean FD vs Variance explained for the reduced (low motion) cohorts for UCLA, Beijing and Cambridge cohorts (See main text). $R^2$ values and the corresponding denoising pipelines are labelled on each panel.

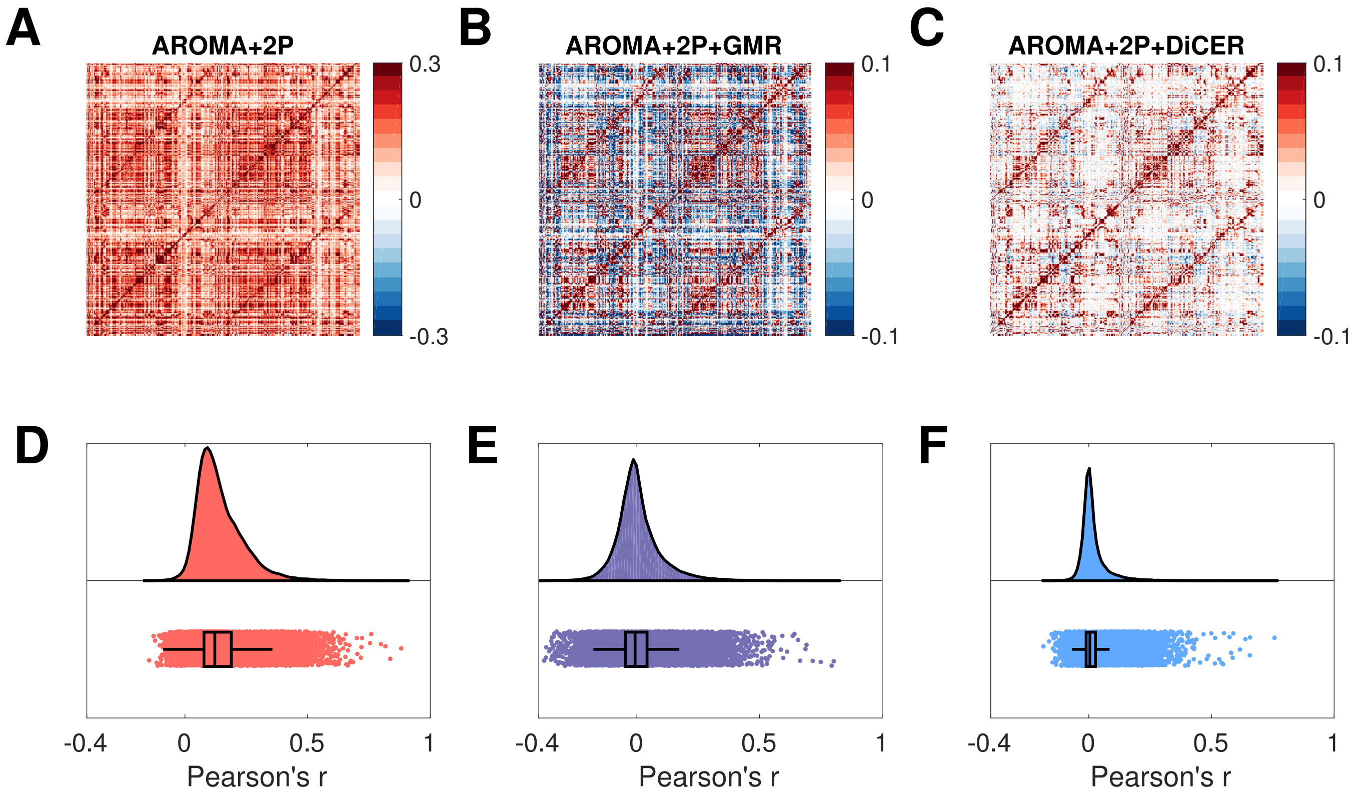


**Fig. S12. Functional connectivity (FC) depends strongly on fMRI preprocessing.** We plot FC estimates for: **A** ICA-AROMA+2P, **B** ICA-AROMA+2P+GMR, and **C** ICA-AROMA+2P+DiCER. FC was estimated as the Pearson correlations between the mean signal in each pair of regions for a 333-ROI-per-hemisphere cortical parcellation (3). The group-level estimates shown here were computed by edge-wise averaging of the $z$-transformed FC matrices over all Beijing subjects. Nodes are ordered in the left hemisphere (upper half) followed by the right hemisphere (lower half). Note that the color bar range varies between plots: **A**: $\pm 0.3$, **B**–**C**: $\pm 0.1$. Beneath each panel **A**–**C** i.e., **D**–**F**; are distributions of the correlation maps (Pearson's $r$ of the upper triangle of the FC matrices) as smoothed kernel density estimates at top and boxplots at bottom, with medians and interquartile ranges annotated (2).

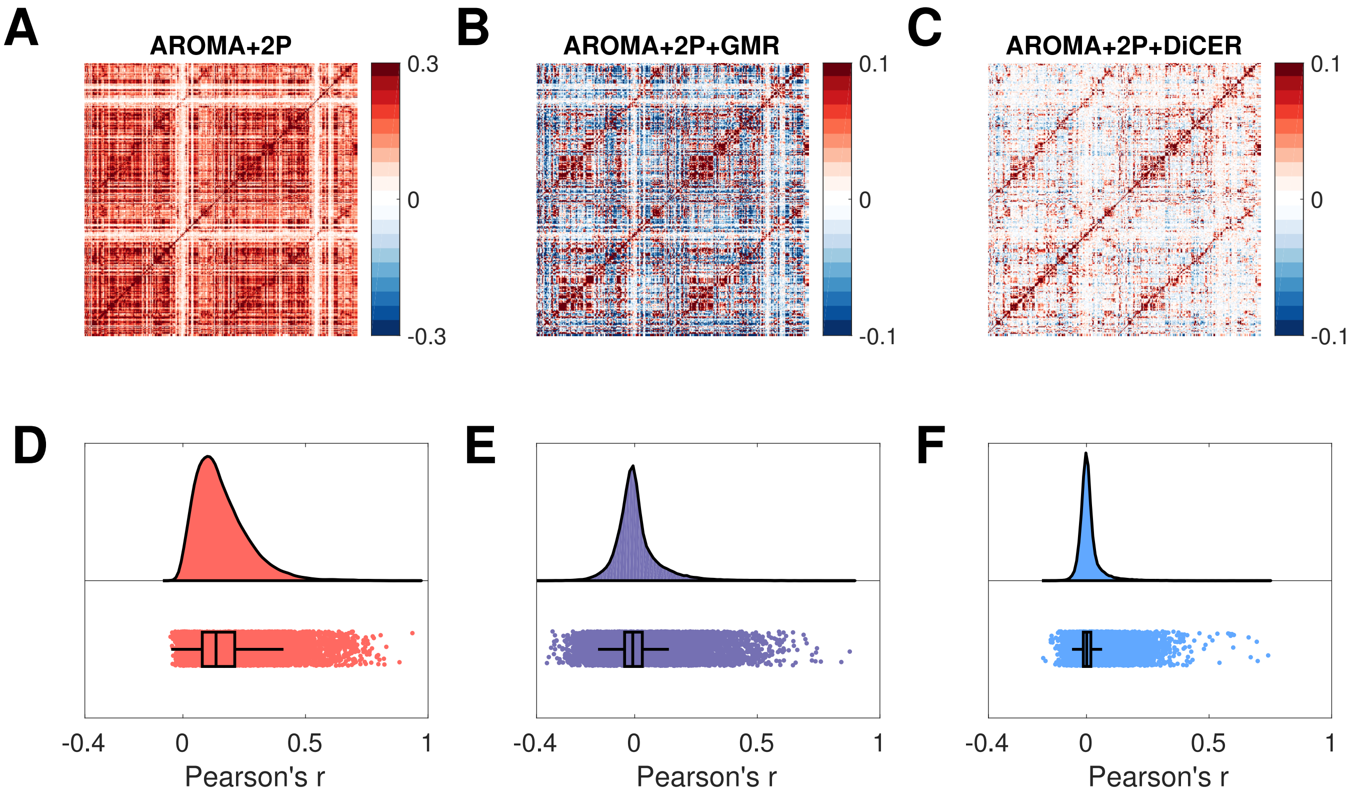


**Fig. S13. Functional connectivity (FC) depends strongly on fMRI preprocessing.** We plot FC estimates for: **A** ICA-AROMA+2P, **B** ICA-AROMA+2P+GMR, and **C** ICA-AROMA+2P+DiCER. FC was estimated as the Pearson correlations between the mean signal in each pair of regions for a 333-ROI-per-hemisphere cortical parcellation (3). The group-level estimates shown here were computed by edge-wise averaging of the $z$-transformed FC matrices over all Cambridge subjects. Nodes are ordered in the left hemisphere (upper half) followed by the right hemisphere (lower half). Note that the color bar range varies between plots: **A**: $\pm 0.3$, **B**–**C**: $\pm 0.1$. Beneath each panel **A**–**C** i.e., **D**–**F**; are distributions of the correlation maps (Pearson's $r$ of the upper triangle of the FC matrices) as smoothed kernel density estimates at top and boxplots at bottom, with medians and interquartile ranges annotated (2).
