## Supplementary Text 2 for "Identifying and removing widespread signal deflections from fMRI data: Rethinking the global signal regression problem"

### Supplementary Text 2 (IC Components): ‘Identifying and removing widespread signal deflections in fMRI data: Rethinking the global signal regression problem’.

**Kevin M. Aquino**[1,*], **Ben D. Fulcher**[2], **Linden Parkes**[1], **Kristina Sabaroedin**[1], **and Alex Fornito**[1]

[1]The Turner Institute for Brain and Mental Health, School of Psychological Sciences, and Monash Biomedical Imaging, Monash University, Victoria, Australia.
[2]School of Physics, The University of Sydney, NSW 2006, Australia.

| IC component | ICA-AROMA+2P | ICA-AROMA+2P +GMR | ICA-AROMA+2P+DiCER |
|---|---|---|---|
| 1 | noise | noise | **pCC/pDMN** |
| 2 | noise | noise | **Visual** |
| 3 | noise | noise | **Medial DMN** |
| 4 | noise | noise | **SMC** |
| 5 | **CON** | noise | **left FPN** |
| 6 | noise | noise | **Hippocampal** |
| 7 | noise | **pDMN** | **left FPN** |
| 8 | **Hippocampal** | noise | **DAN** |
| 9 | **pDMN** | **Hippocampal** | **right FPN** |
| 10 | Visual | **LFPN** | **DAN** |

**Table S1. The first ten MELODIC components are concentrated in meaningful functional networks when using DiCER for the Beijing dataset**. In each denoising stream, bold font indicates a spatial IC component that corresponds to a classic resting-state network (1).

#### Independent components and dual regression results for: UCLA

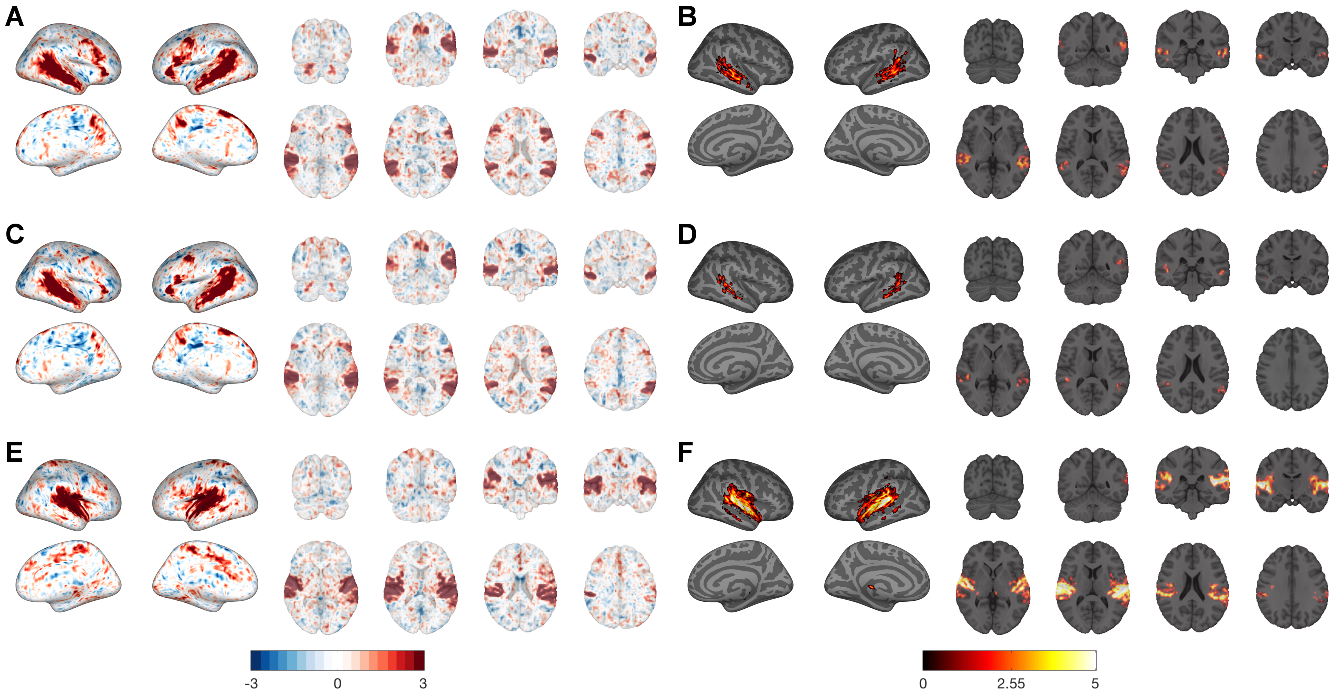


**Fig. S13.** Auditory resting-state network uncovered using melodic within the UCLA Cohort. The group-level independent component maps, with the corresponding activation map for AROMA+2P, AROMA+2P+GMR, and AROMA+2P+DiCER are shown in the pairs **A,B**, **C,D**, and **E,F** and respectively. The group-level activation maps are calculated from dual-regression, and the thresholded t-statistics are displayed.

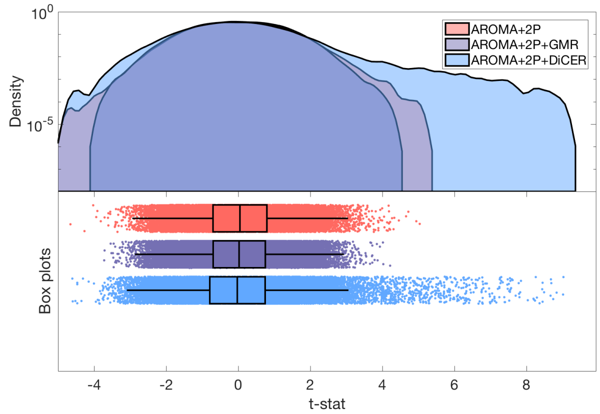


**Fig. S14.** The full distribution for the $t$-statistic at the group-level for the IC component that represents the Auditory resting-state network within the UCLA Cohort for all three de-noising pipelines. The distributions are shown as smoothed kernel density estimates at top and boxplots at bottom, with medians and interquartile ranges annotated (2). Note the logarithmic vertical axis for this kernel density to resolve the tails of this distribution.

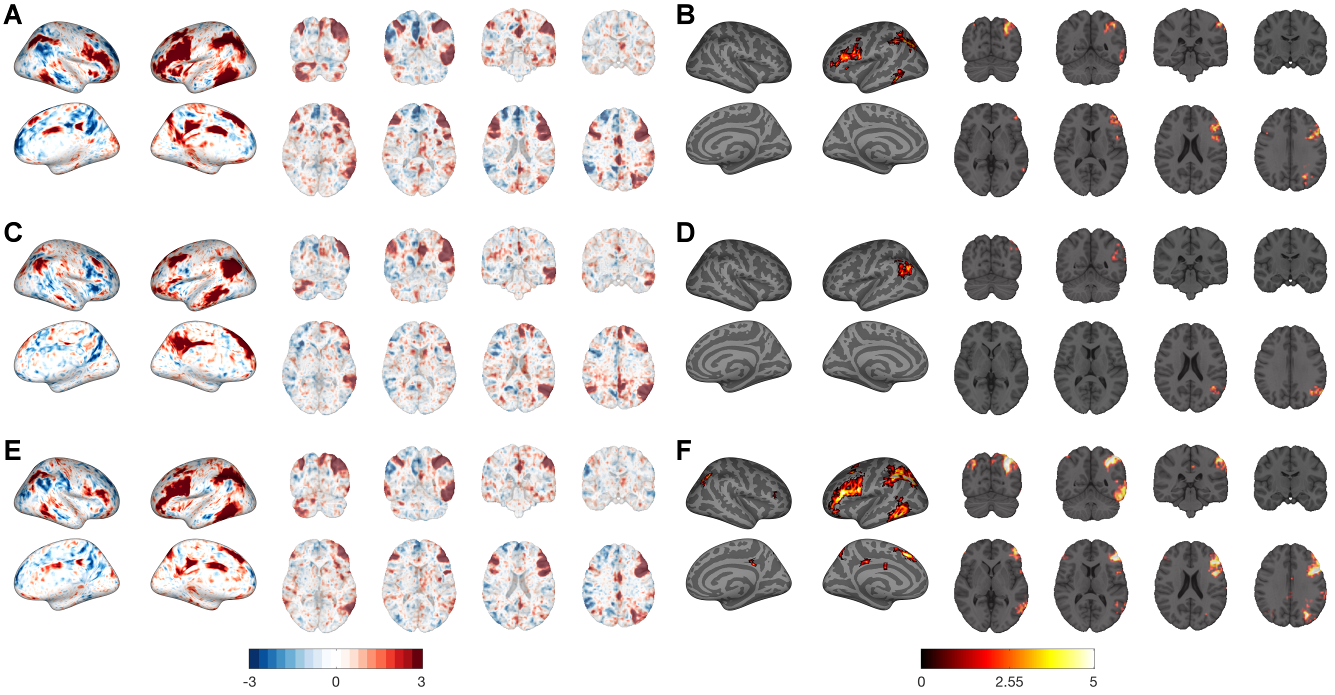


**Fig. S15.** Left FPN resting-state network uncovered using melodic within the UCLA Cohort. The group-level independent component maps, with the corresponding activation map for AROMA+2P, AROMA+2P+GMR, and AROMA+2P+DiCER are shown in the pairs **A,B**, **C,D**, and **E,F** and respectively. The group-level activation maps are calculated from dual-regression, and the thresholded t-statistics are displayed.

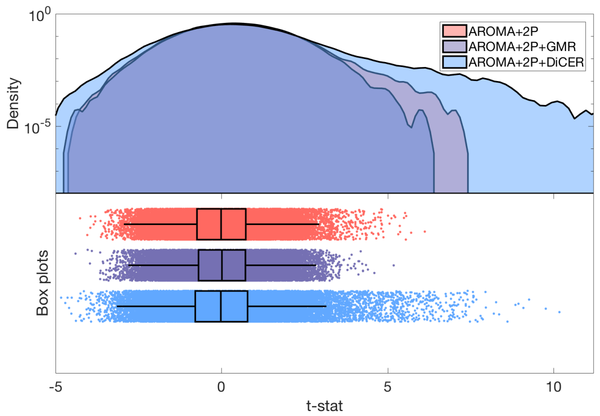


**Fig. S16.** The full distribution for the $t$-statistic at the group-level for the IC component that represents the Left FPN resting-state network within the UCLA Cohort for all three de-noising pipelines. The distributions are shown as smoothed kernel density estimates at top and boxplots at bottom, with medians and interquartile ranges annotated (2). Note the logarithmic vertical axis for this kernel density to resolve the tails of this distribution.

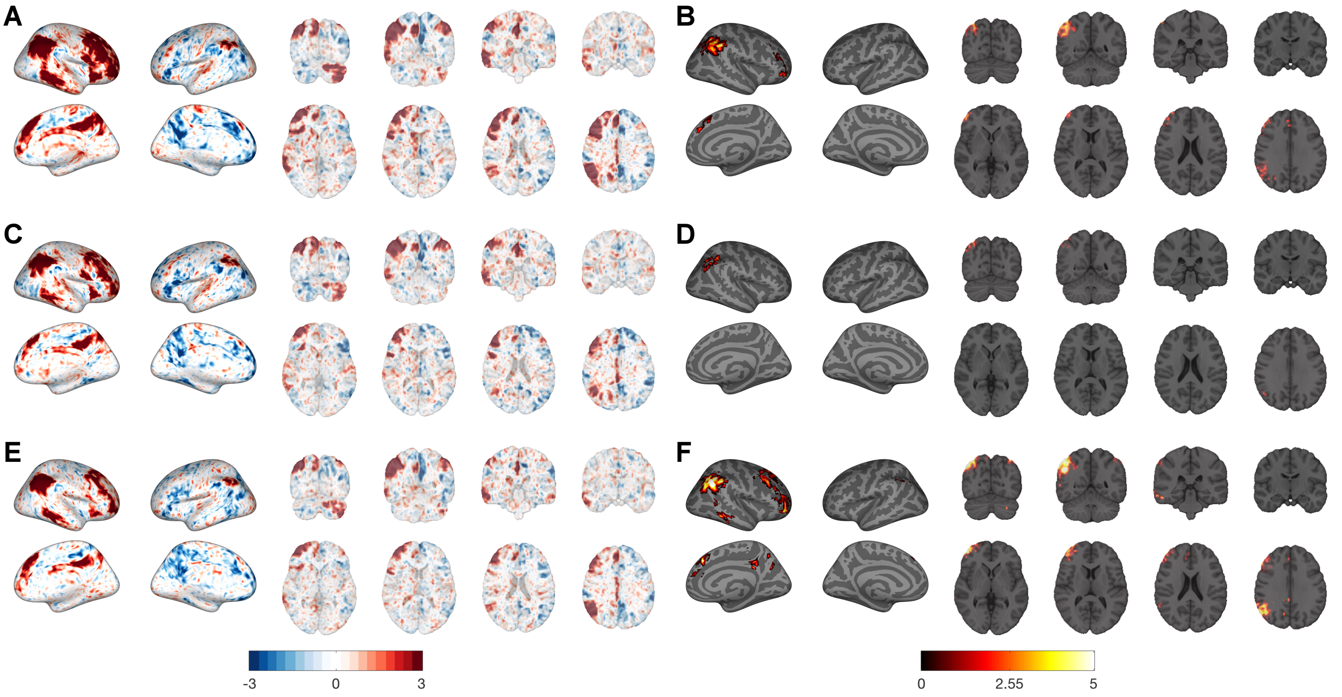


**Fig. S17.** Right FPN resting-state network uncovered using melodic within the UCLA Cohort. The group-level independent component maps, with the corresponding activation map for AROMA+2P, AROMA+2P+GMR, and AROMA+2P+DiCER are shown in the pairs **A,B**, **C,D**, and **E,F** and respectively. The group-level activation maps are calculated from dual-regression, and the thresholded $t$-statistics are displayed.

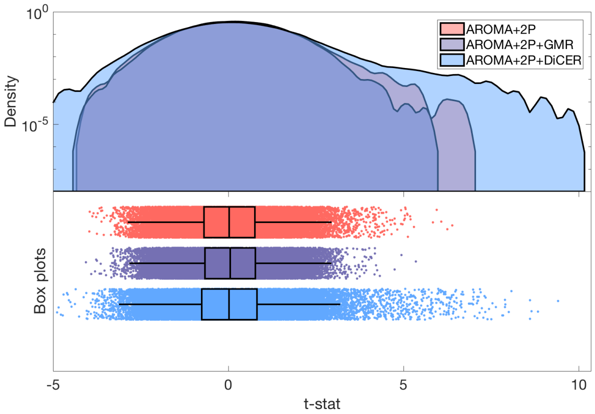


**Fig. S18.** The full distribution for the $t$-statistic at the group-level for the IC component that represents the Right FPN resting-state network within the UCLA Cohort for all three de-noising pipelines. The distributions are shown as smoothed kernel density estimates at top and boxplots at bottom, with medians and interquartile ranges annotated (2). Note the logarithmic vertical axis for this kernel density to resolve the tails of this distribution.

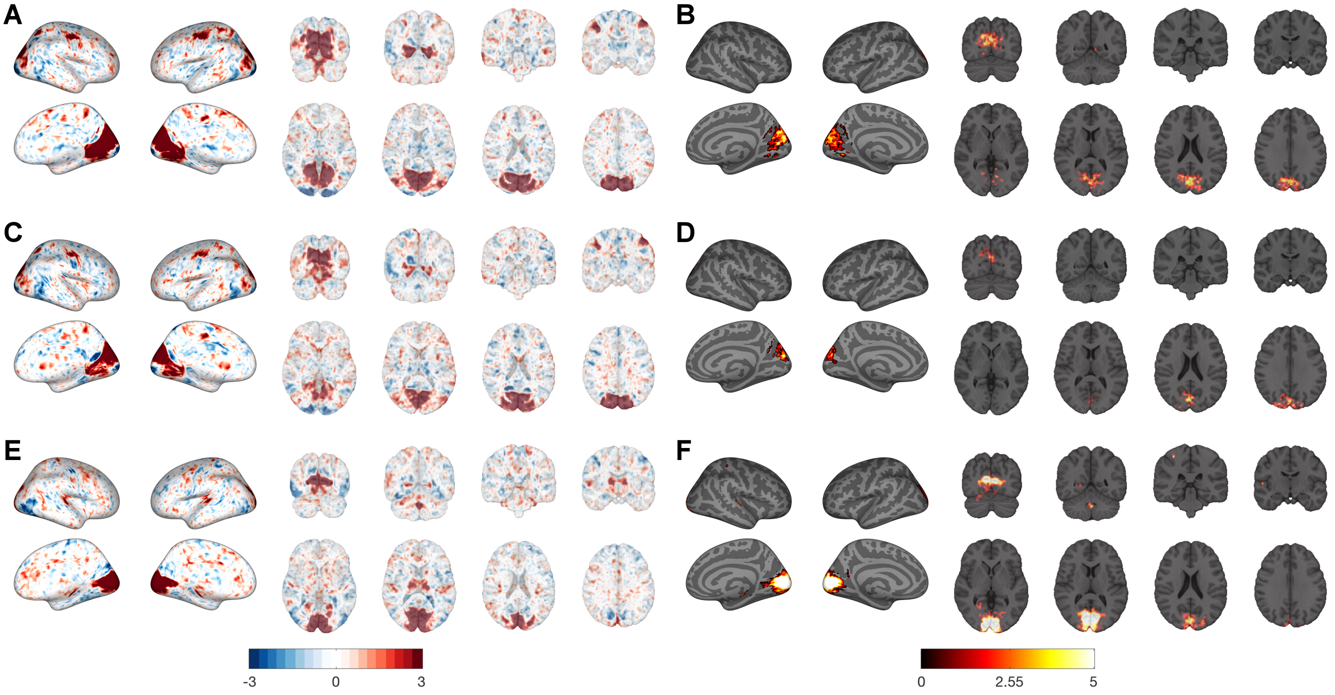


**Fig. S19.** Visual resting-state network uncovered using melodic within the UCLA Cohort. The group-level independent component maps, with the corresponding activation map for AROMA+2P, AROMA+2P+GMR, and AROMA+2P+DiCER are shown in the pairs **A,B**, **C,D**, and **E,F** and respectively. The group-level activation maps are calculated from dual-regression, and the thresholded $t$-statistics are displayed.

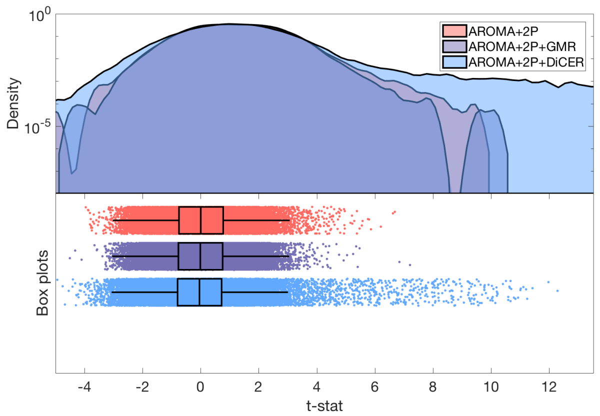


**Fig. S20.** The full distribution for the $t$-statistic at the group-level for the IC component that represents the Visual resting-state network within the UCLA Cohort for all three de-noising pipelines. The distributions are shown as smoothed kernel density estimates at top and boxplots at bottom, with medians and interquartile ranges annotated (2). Note the logarithmic vertical axis for this kernel density to resolve the tails of this distribution.

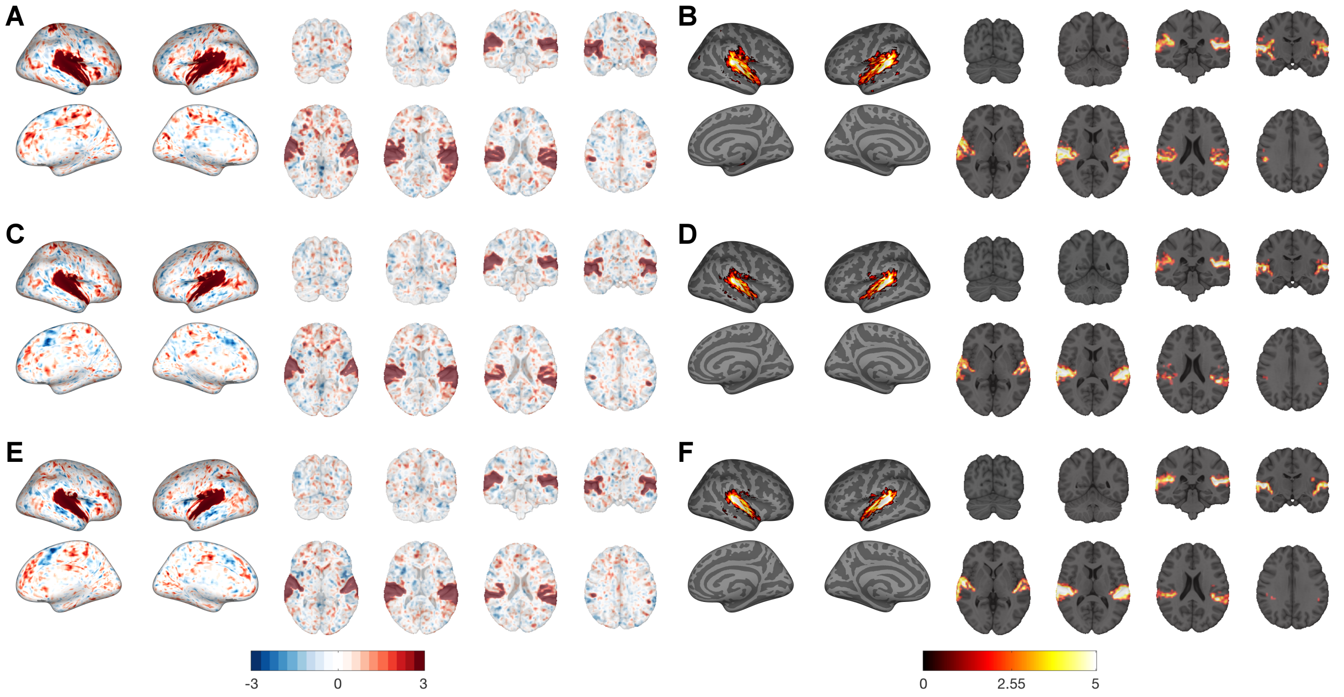


**Fig. S21.** Auditory resting-state network (AUD) uncovered using melodic within the Beijing Cohort. The group-level independent component maps, with the corresponding activation map for AROMA+2P, AROMA+2P+GMR, and AROMA+2P+DiCER are shown in the pairs **A,B**, **C,D**, and **E,F** and respectively. The group-level activation maps are calculated from dual-regression, and the thresholded $t$-statistics are displayed.

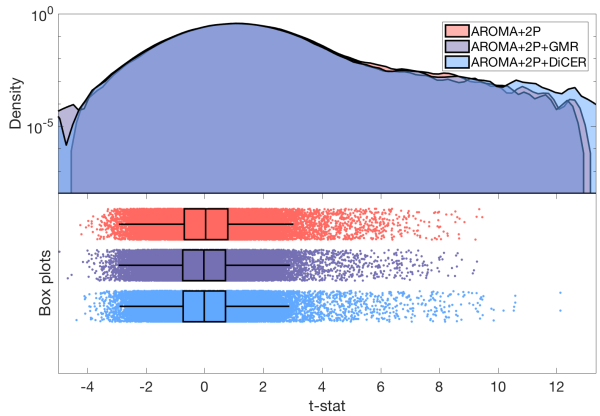


**Fig. S22.** The full distribution for the $t$-statistic at the group-level for the IC component that represents the AUD within the Beijing Cohort for all three de-noising pipelines. The distributions are shown as smoothed kernel density estimates at top and boxplots at bottom, with medians and interquartile ranges annotated (2). Note the logarithmic vertical axis for this kernel density to resolve the tails of this distribution.

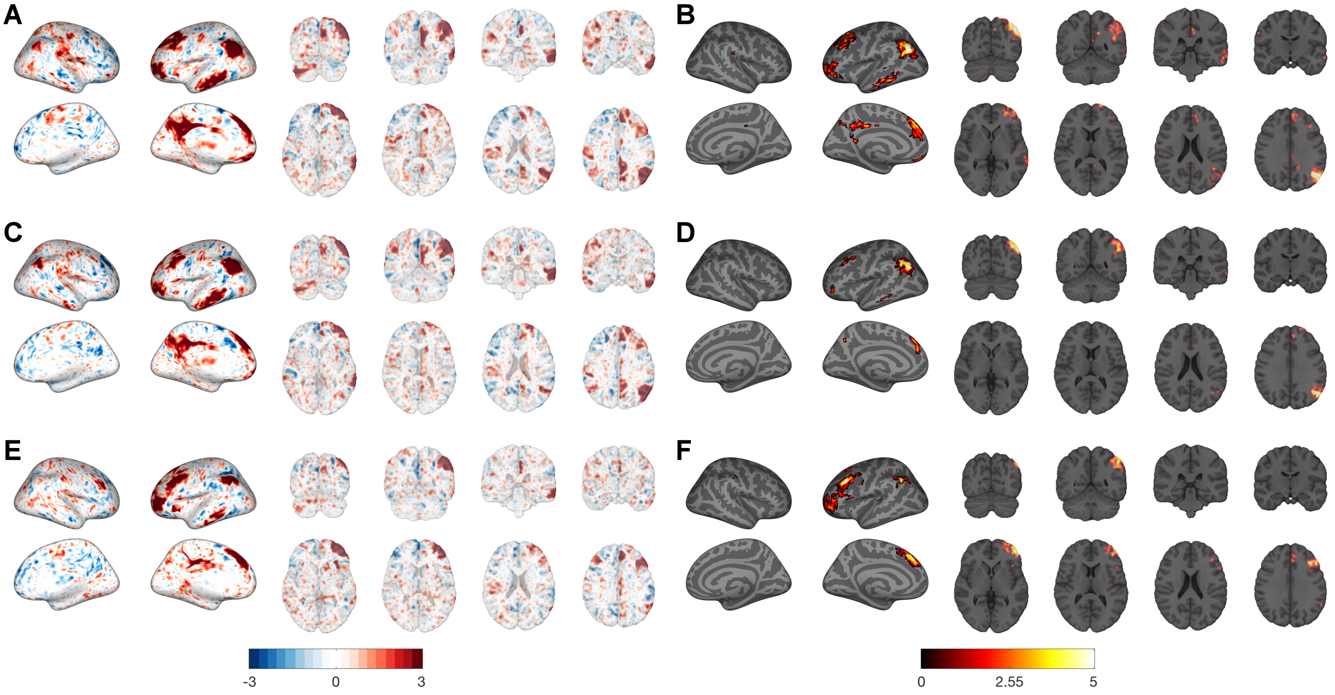


**Fig. S23.** Left FPN resting-state network uncovered using melodic within the Beijing Cohort. The group-level independent component maps, with the corresponding activation map for AROMA+2P, AROMA+2P+GMR, and AROMA+2P+DiCER are shown in the pairs **A,B**, **C,D**, and **E,F** and respectively. The group-level activation maps are calculated from dual-regression, and the thresholded $t$-statistics are displayed.

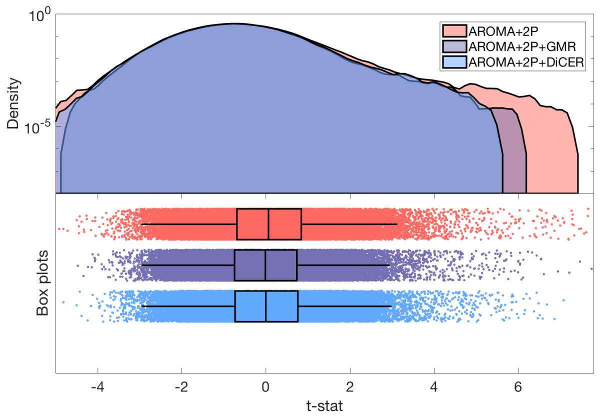


**Fig. S24.** The full distribution for the $t$-statistic at the group-level for the IC component that represents the LFPN within the Beijing Cohort for all three de-noising pipelines. The distributions are shown as smoothed kernel density estimates at top and boxplots at bottom, with medians and interquartile ranges annotated (2). Note the logarithmic vertical axis for this kernel density to resolve the tails of this distribution.

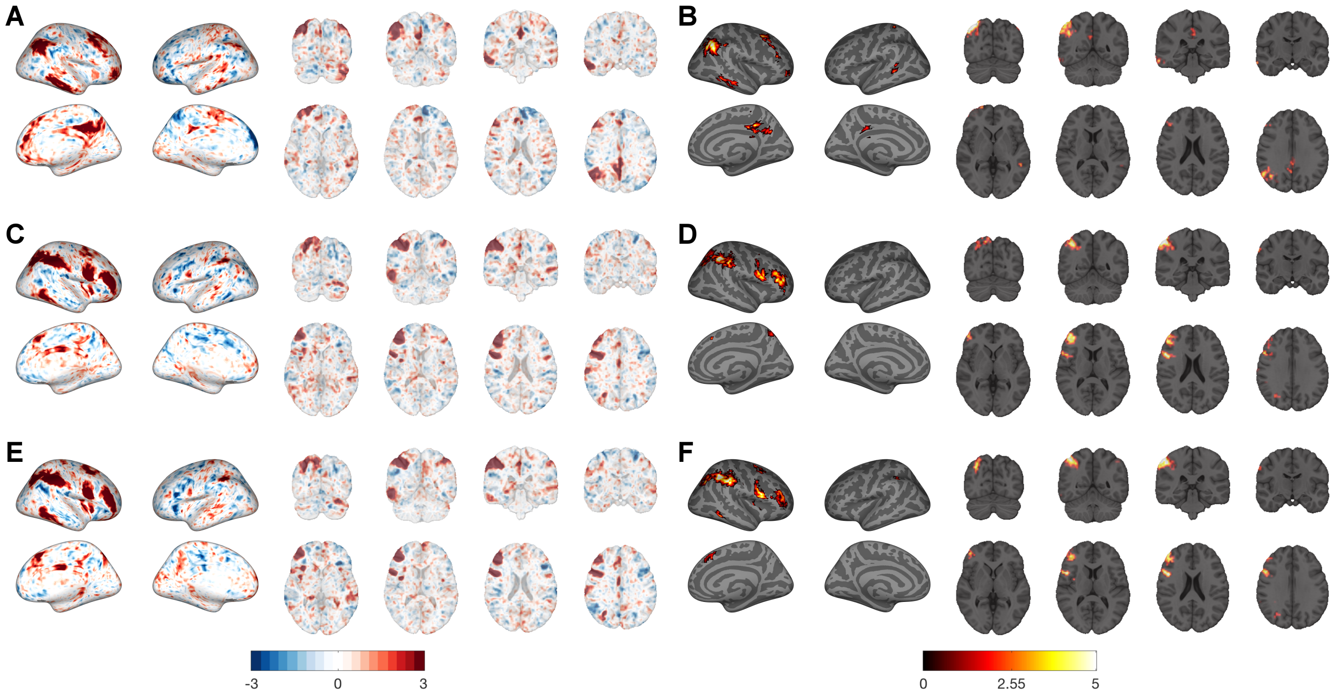


**Fig. S25.** Right FPN resting-state network uncovered using melodic within the Beijing Cohort. The group-level independent component maps, with the corresponding activation map for AROMA+2P, AROMA+2P+GMR, and AROMA+2P+DiCER are shown in the pairs **A,B**, **C,D**, and **E,F** and respectively. The group-level activation maps are calculated from dual-regression, and the thresholded $t$-statistics are displayed.

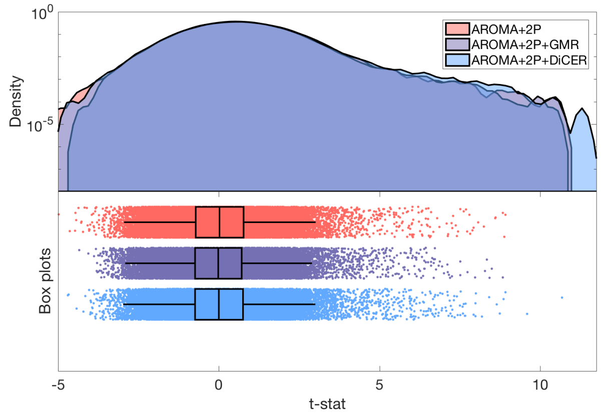


**Fig. S26.** The full distribution for the $t$-statistic at the group-level for the IC component that represents the RFPN within the Beijing Cohort for all three de-noising pipelines. The distributions are shown as smoothed kernel density estimates at top and boxplots at bottom, with medians and interquartile ranges annotated (2). Note the logarithmic vertical axis for this kernel density to resolve the tails of this distribution.

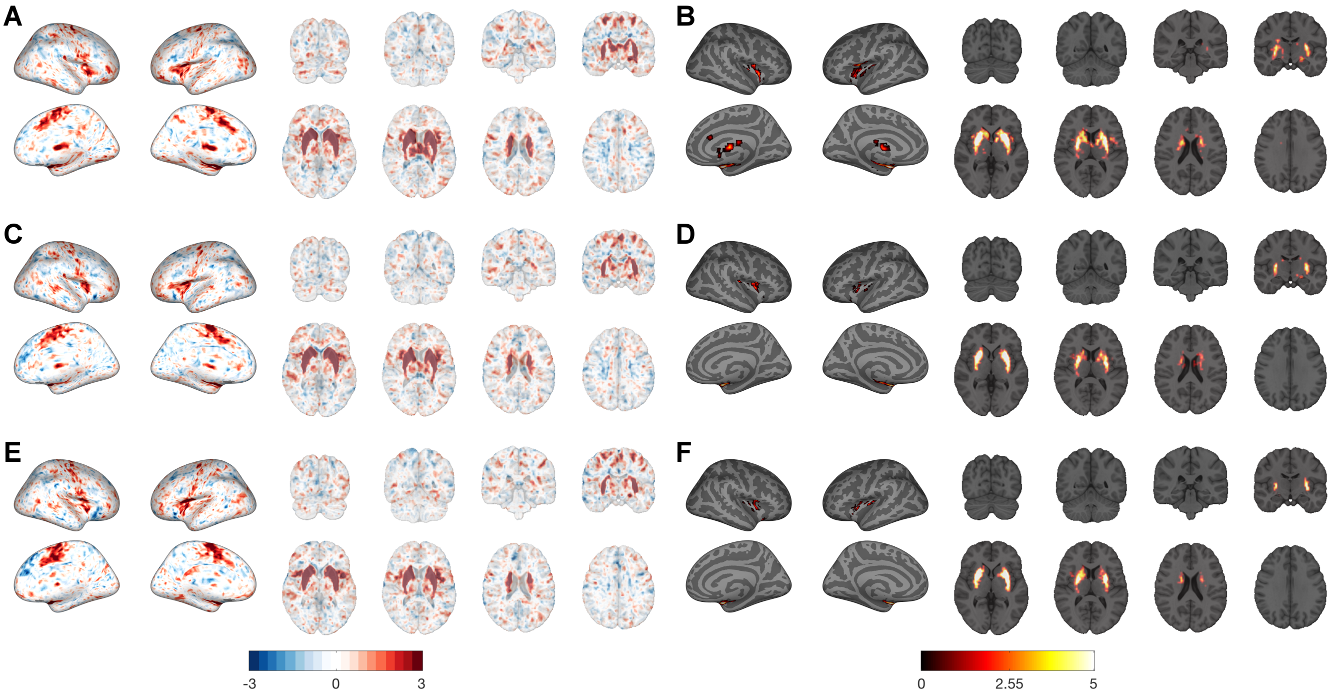


**Fig. S27.** Putamen resting-state network uncovered using melodic within the Beijing Cohort. The group-level independent component maps, with the corresponding activation map for AROMA+2P, AROMA+2P+GMR, and AROMA+2P+DiCER are shown in the pairs **A,B**, **C,D**, and **E,F** and respectively. The group-level activation maps are calculated from dual-regression, and the thresholded $t$-statistics are displayed.

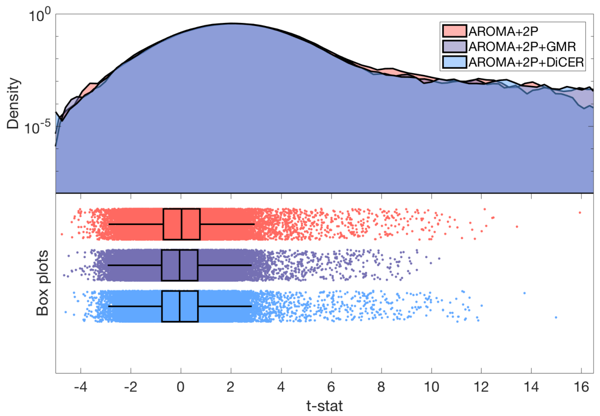


**Fig. S28.** The full distribution for the $t$-statistic at the group-level for the IC component that represents the Striatum+Putamen within the Beijing Cohort for all three de-noising pipelines. The distributions are shown as smoothed kernel density estimates at top and boxplots at bottom, with medians and interquartile ranges annotated (2). Note the logarithmic vertical axis for this kernel density to resolve the tails of this distribution.

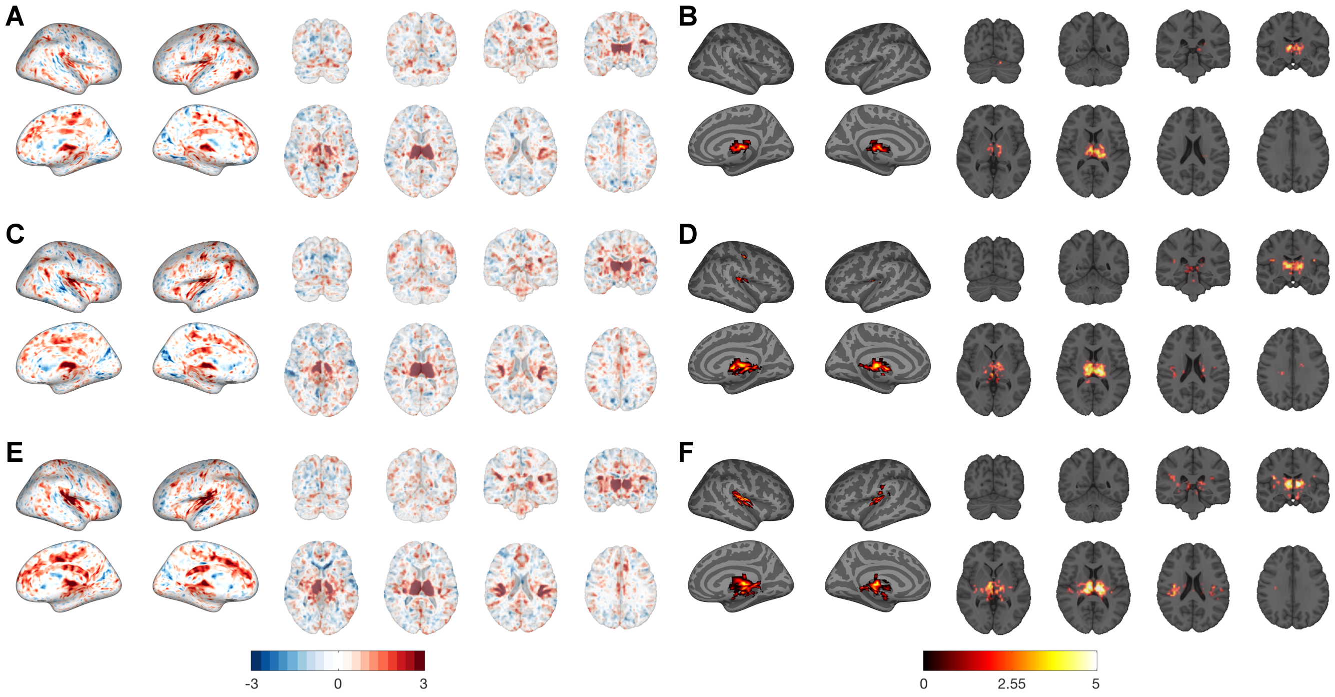


**Fig. S29.** Thalamic resting-state network uncovered using melodic within the Beijing Cohort. The group-level independent component maps, with the corresponding activation map for AROMA+2P, AROMA+2P+GMR, and AROMA+2P+DiCER are shown in the pairs **A,B**, **C,D**, and **E,F** and respectively. The group-level activation maps are calculated from dual-regression, and the thresholded $t$-statistics are displayed.

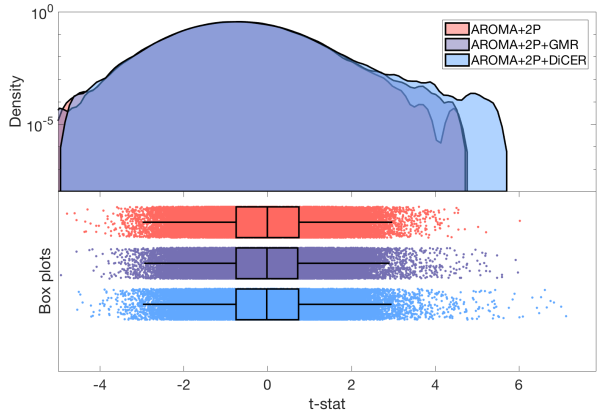


**Fig. S30.** The full distribution for the $t$-statistic at the group-level for the IC component that represents the Thalamus within the Beijing Cohort for all three de-noising pipelines. The distributions are shown as smoothed kernel density estimates at top and boxplots at bottom, with medians and interquartile ranges annotated (2). Note the logarithmic vertical axis for this kernel density to resolve the tails of this distribution.

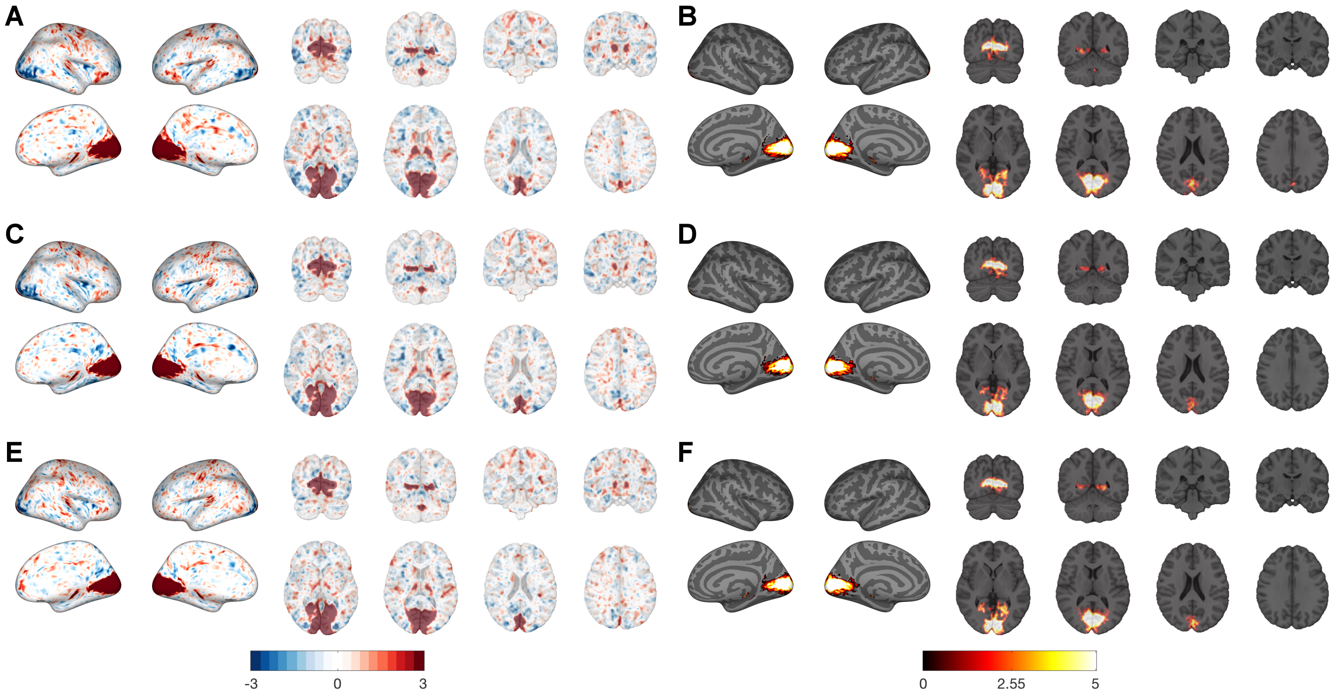


**Fig. S31.** Visual resting-state network uncovered using melodic within the Beijing Cohort. The group-level independent component maps, with the corresponding activation map for AROMA+2P, AROMA+2P+GMR, and AROMA+2P+DiCER are shown in the pairs **A,B**, **C,D**, and **E,F** and respectively. The group-level activation maps are calculated from dual-regression, and the thresholded $t$-statistics are displayed.

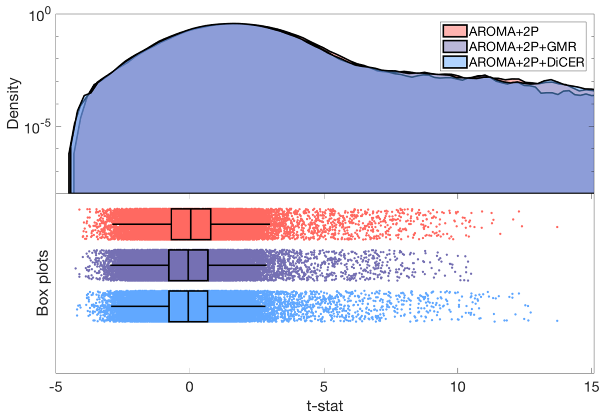


**Fig. S32.** The full distribution for the $t$-statistic at the group-level for the IC component that represents the Visual resting-state network within the Beijing Cohort for all three de-noising pipelines. The distributions are shown as smoothed kernel density estimates at top and boxplots at bottom, with medians and interquartile ranges annotated (2). Note the logarithmic vertical axis for this kernel density to resolve the tails of this distribution.
